# Target-enriched enzymatic methyl sequencing: flexible, scalable and inexpensive hybridization capture for quantifying DNA methylation

**DOI:** 10.1101/2022.08.26.505457

**Authors:** Dustin R. Rubenstein, Joseph Solomon

## Abstract

The increasing interest in studying DNA methylation to understand how traits or diseases develop requires new and flexible approaches for quantifying DNA methylation in a diversity of organisms. In particular, we need efficient yet cost-effective ways to measure CpG methylation states over large and complete regions of the genome. Here, we develop TEEM-Seq (target-enriched enzymatic methyl sequencing), a method that combines enzymatic methyl sequencing with a custom-designed hybridization capture bait set that can be scaled to reactions including large numbers of samples in any species for which a reference genome is available. Using DNA from a passerine bird, the superb starling (*Lamprotornis superbus*), we show that TEEM-Seq is able to quantify DNA methylation states similarly well to the more traditional approaches of whole-genome and reduced-representation sequencing. Moreover, we demonstrate its reliability and repeatability, as duplicate libraries from the same samples were highly correlated. Importantly, the downstream bioinformatic analysis for TEEM-Seq is the same as for any sequence-based approach to studying DNA methylation, making it simple to incorporate into a variety of workflows. We believe that TEEM-Seq could replace traditional approaches for studying DNA methylation in candidate genes and pathways, and be effectively paired with other whole-genome or reduced-representation sequencing approaches to increase project sample sizes. In addition, TEEM-Seq can be combined with mRNA sequencing to examine how DNA methylation in promoters or other regulatory regions is related to the expression of individual genes or gene networks. By maximizing the number of samples in the hybridization reaction, TEEM-Seq is an inexpensive and flexible sequence-based approach for quantifying DNA methylation in species where other capture-based methods are unavailable or too expensive, particularly for non-model organisms.

## Introduction

There is increasing recognition in fields as diverse as ecology, evolutionary biology, conservation biology, neuroscience, and biomedicine that epigenetics is critical to understanding how traits, diseases, or other phenotypes develop [1–4]. One of the most widely studied epigenetic mechanisms underlying phenotypic variation is DNA methylation [5,6], the addition of a methyl group to a DNA nucleotide that often impacts gene expression through transcriptional silencing [7]. In mammals and many other vertebrates, DNA methylation typically occurs at CpG sites, or regions of DNA where a cytosine nucleotide is followed by a guanine nucleotide. With the increasing availability of reference genomes for non-model organisms and the continually decreasing costs of high-throughput sequencing, sequencing-based approaches have become the preferred method for studying DNA methylation across the diversity of life because they can provide quantitative values of CpG methylation states [8]. Sequence-based approaches to studying DNA methylation range from whole- or reduced-representation sequencing of genomes to sequencing one or a few candidate genes, yet the performance and cost of different sequence-based approaches to quantify and map DNA methylation varies greatly [9–12]. Whole-genome sequencing for DNA methylation profiling remains the gold standard, but is costly because of the necessary sequencing depth, particularly for species with large genomes [8]. In contrast, reduced-representation approaches are cheaper than whole-genome sequencing because they cover only a fraction of the genome, but since such approaches require restriction enzymes to target specific genomic regions (e.g., CpG islands), they inevitably miss potentially important CpG sites in genomic regions of interest [13]. Despite the differences in cost and coverage, most whole-genome and reduced-representation sequencing approaches share the chemical conversion of unmethylated cytosines to uracils with bisulfite treatment [14], a method that enables quantification of methylation states but that can fragment DNA and complicate downstream library preparation and even bias results [12,15].

As the number of studies of DNA methylation grows, it becomes increasingly important to develop new methods that can quantitatively measure CpG methylation states over large and complete regions of the genome in an efficient yet cost-effective way. Such approaches exist for humans and some model organisms in the form of bead or probe panels, but flexible methods for studying targeted complete genomic regions of varying size in any organism are much rarer. Although hybridization capture using biotinylated oligonucleotide baits (probes) to hybridize specific regions of interest across the genome of any species have become standard in molecular biology [16], their application in epigenetics has been more limited [17–19]. Commercial approaches such as Agilent’s SureSelect^XT^ Methyl-Seq Target Enrichment System and Roche’s SeqCap Epi Enrichment System are neither particularly scalable (i.e., limited in the number of samples per reaction) nor cost-effective, and both use bisulfite conversion. Yet, with the recent development of alternative approaches to bisulfite treatment for converting DNA [20], newer methods of hybridization capture may be viable for quantitatively studying DNA methylation, particularly in non-model organisms.

Here, we develop a flexible, scalable, and inexpensive method called target-enriched enzymatic methyl sequencing (TEEM-Seq) that uses hybridization capture to quantitively measure CpG methylation states. We achieve this goal by combining enzymatic methyl sequencing (EM-seq) [20] with a custom-designed, species-specific hybridization capture bait set that can be scaled to reactions that include much larger numbers of samples than other currently available methods and that only require low amounts of input DNA to quantify both 5mC and 5hmC methylation. EM-seq effectively overcomes DNA fragmentation caused by bisulfite treatment, and a custom bait set allows for cost-effective and efficient targeting of complete genomic regions. In this study, we selected genomic regions encompassing selected genes known to be associated with the hypothalamic–pituitary–adrenal axis (HPA), the hypothalamic–pituitary–gonadal axis (HPG), and DNA methyltransferase activity. We chose these specific HPA and HPG genes because of the critical roles that they play in the vertebrate stress response and in reproduction [21]. After designing baits targeting all of the putative promoters (roughly 4 kb upstream of the transcription start site) and some of the exons of 22 of these genes, we developed the TEEM-Seq protocol using DNA extracted from the blood of the superb starling (*Lamprotornis superbus*), a passerine bird with a published reference genome [22]. To gauge the effectiveness and efficiency of TEEM-Seq against more traditional methods of quantifying DNA methylation, we compared data from some of the same individual starlings to data from whole-genome enzymatic methyl sequencing (WGEM-Seq) and/or reduced-representation bisulfite sequencing (RRBS) (Fig 1). We found that necessary sequencing coverage levels were similar to RRBS, but an order of magnitude lower than for WGEM-Seq for similarly high-quality coverage of complete target regions. With sequencing depth needs similar to that of reduced-representation approaches, but with complete coverage of genomic regions more similar to results from whole-genome approaches, TEEM-Seq is a cost-effective approach for targeting specific but complete parts of the genome. Ultimately, we demonstrate the utility of TEEM-Seq for quantifying DNA methylation in an efficient and adaptable manner that should be applicable to studying any organism in a variety of tissue and sample types.

**Fig 1.**
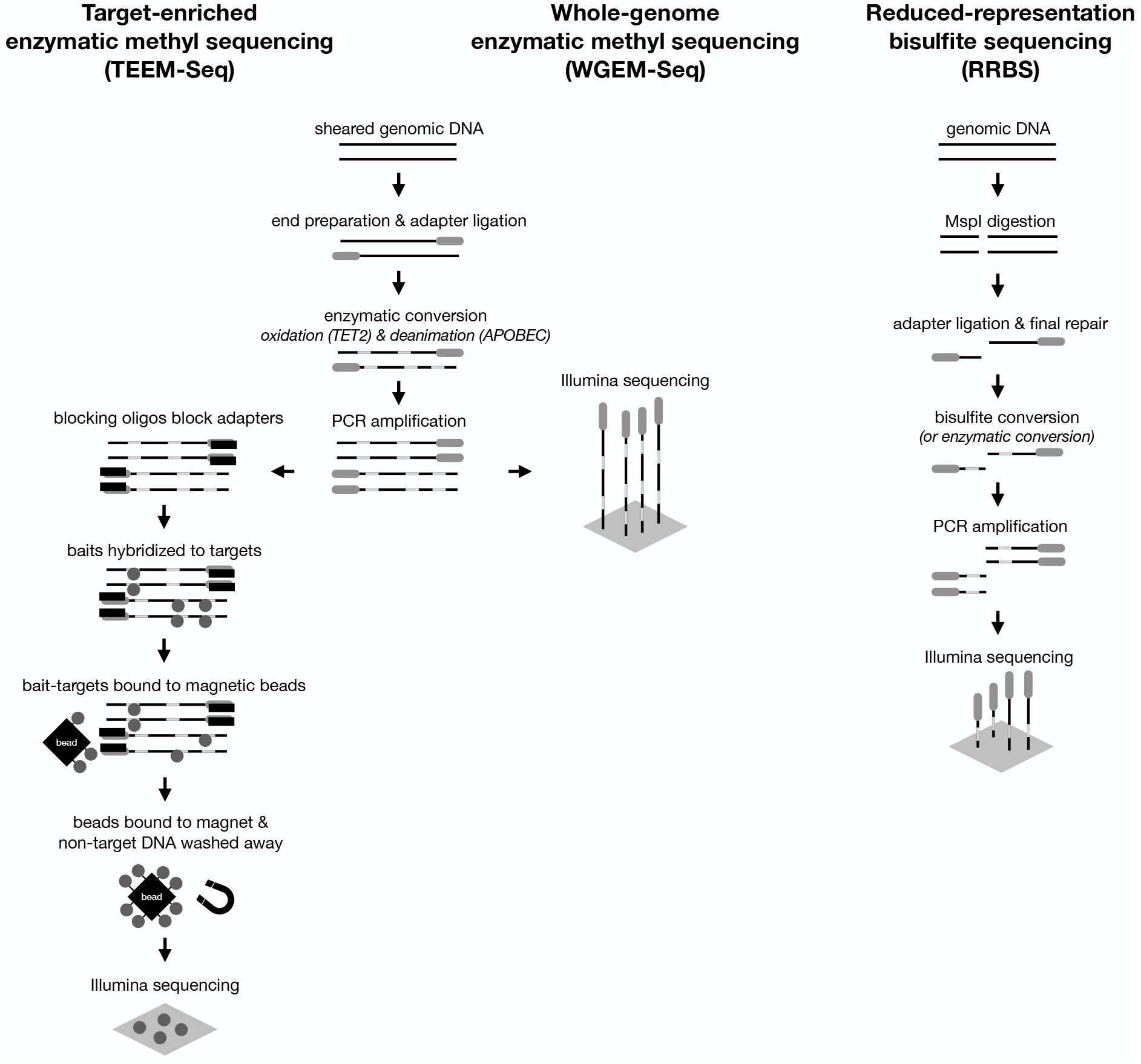
Diagram comparing the workflow for target-enriched enzymatic methyl sequencing (TEEM-Seq), whole-genome enzymatic methyl sequencing (WGEM-Seq), and reduced-representation bisulfite sequencing (RRBS).

## Materials and methods

### Sample collection

Superb starlings (N = 39) were captured using baited traps or mist nests at the Mpala Research Centre, Laikipia, Kenya (0° 17’N, 37° 52’E). Whole blood was collected in 2% SDS Queens lysis buffer [23] and genomic DNA was extracted using a DNeasy Blood & Tissue Kit (Qiagen). All research was approved by the Institutional Animal Care and Use Committee at Columbia University (AC-AAAW6451), as well as the Kenyan National Commission for Science, Technology and Innovation, the Kenyan National Environmental Management Authority, the Kenya Wildlife Service, the Kenyan Wildlife Research and Training Institute, and the Mpala Research Centre.

### Library preparation

After DNA was quantified using a Qubit 3.0 Fluorometer (Thermo Fisher Scientific), 100-150 ng was sheared to 250-275 bp on a Covaris ME220 and inspected on an Agilent 4200 TapeStation. The sheared DNA was then used as a template for libraries prepared with a NEBNext Enzymatic Methyl-seq Kit (EM-seq) (New England Biolabs). We note that our previous attempts to develop a hybridization capture protocol using bisulfite-converted DNA were unsuccessful, presumably because chemical conversion led to fragmented DNA that failed in hybridization reactions. However, the enzymatic conversion overcame challenges associated with fragmented DNA from bisulfite conversion and allowed for successful hybridization capture. Following the EM-seq Kit protocol, sheared DNA was end repaired at 20°C, ligated with an EM-seq adaptor at 20°C, and then used as template in a TET2 reaction at 37°C for oxidation of 5-Methylcytosines and 5-Hydroxymethylcytosines. The TET2-converted DNA was denatured with formamide at 85°C, APOBEC cytosine deaminated at 37°C, and then PCR amplified with index primers and NEBNext Q5U Master Mix at an annealing temperature of 62°C. Since we intended to perform downstream in-solution hybridization capture, we increased library yield through 12 total PCR cycles for the enzymatic conversion amplification compared with the standard recommendation of 4-8 cycles. Although we did not vary the number of PCR cycles in this study, this could be optimized in the future to possibly reduce PCR duplicates. Whereas the duplicate rate increased most in the hybridization reaction and amplification, it remains unknown whether EM-seq duplicates were compounded in the capture process through bait bias toward more highly represented duplicate molecules. We also increased all SPRI bead volumes by 50%, supplementing the included sample purification beads with AMPure XP beads (Beckman Coulter) at the PCR clean-up stage, because we intended the EM-seq libraries for a hybridization capture pool rather than for direct sequencing, where free adaptor carryover could be an issue. A 50% increase was chosen to improve library yield without excessive carryover of free adaptors and at a modest elevation in clean-up cost. Although we did not vary the ratios of beads to sample, this too could be further optimized to reduce costs. For post-ligation clean-up, we used a ratio of beads to sample volume of 1.76:1 instead of the NEB EM-seq standard protocol’s 1.18:1; for the TET2 reaction cleanup, we used a ratio of 2.65:1 instead of 1.76:1; for the APOBEC deamination clean-up we used 1.5:1 instead of 1:1; and for post-amplification clean-up we used 1.35:1 instead of 0.9:1. In addition, we varied kit reagent volumes down to half of those recommended, ultimately settling on three quarter volumes for our final reactions as a way to reduce costs. Results with three quarter volumes of reagents worked similarly well to those with full volumes. Finally, we Qubit-quantified the libraries, inspected them on the TapeStation, and pooled 25 ng from each library into 1.2 μg total for the pool of 48 libraries.

### Bait design

Target sequences for bait design were identified from the superb starling reference genome (CU_Lasu_v1) [22]. Baits targeted 23 regions (*GNRHR2* had two separate regions) of roughly 4 kb in length upstream of the transcription start site (i.e., putative promoter regions) and exons (excluding introns) from a subset of genes known to be associated with the HPA axis, the HPG axis, and DNA methyltransferases (Table 1). Biotinylated RNA baits were then prepared commercially using myBaits v4.01 Custom 1-20K 16 Reaction Kits (Daicel Arbor Biosciences). We submitted 71 sequences (113,940 bp) for 80 nt bait design at 2x tiling density. For each bait candidate, in addition to the unmodified original, myBaits designed eight additional genome variants to reasonably cover conversion permutations (i.e., all methylated, a random 50% CpGs methylated [to account for enzymatic conversion], the other 50% CpGs methylated, unmethylated, and sense/antisense for each version). This resulted in 21,792 baits, all of which were quality-assessed and filtered by myBaits Design based on masked repeats in the original sequences (repeatmasker [engine: cross_match, organism: Aves] resulting in 5.4% of total sequences masked), GC content, and BLAST hits on the zebra finch (*Taeniopygia guttata*), European starling (*Sturnus vulgaris*), collared flycatcher (*Ficedula albicollis*), and great tit (*Parus major*) reference genomes for the final myBaits kits (S1 and S2 Tables). The goal of our multispecies genome-wide BLAST screen was to check for any bait candidates that were likely to be very non-specific in general as seen across any of the four avian genomes (not to filter out any candidates that were blasting to other species’ genomes in conserved regions), which could contribute to low experimental sequencing efficiency by capturing undesired off-target reads. In addition to unmodified versions of each genome, myBaits blasted bait candidates against sense strand unmethylated (i.e., all Cs converted to Ts) and anti-sense strand unmethylated (i.e., all Gs converted to As) versions of each genome, for a total of 12 genome variants. The top 500 blast hits (by bit score) were tabulated for each bait candidate. The final bait set contained 15,308 sequences with <25% repeat-masking, 10-80% GC content, and £10 max BLAST hits across all 12 genome variants to avoid non-specific targets. In other words, starling bait sequences were filtered based on likely performance, including possible off-target capture, and low-complexity baits post-conversion were excluded.

**Table 1.** Targeted genomic regions for superb starling baits. Although we targeted putative promoter regions of all of the genes (i.e., 4 kb upstream of the transcription start site where annotated), we also targeted complete exonic regions (indicated by *) or gene body regions (based on similarity with Ensembl genome sequences, including the collared flycatcher for *GNRH1* and zebra finch for *MC4R*, indicated by **) in a subset of the genes.

| Target region | Gene name |
| --- | --- |
| *AR* * | Androgen receptor |
| *AVPR1A* * | Arginine vasopressin receptor 1A |
| *AVPR1B* * | Arginine vasopressin receptor 1B |
| *CRH* * | Corticotropin releasing hormone |
| *DNMT1* | DNA methyltransferase 1 |
| *DNMT3A* | DNA methyltransferase 3A |
| *DNMT3B* | DNA methyltransferase 3B |
| *EGR1* * | Early growth response 1 |
| *ESR1* | Estrogen receptor 1 |
| *FKBP5* * | FK506 binding protein 5 |
| *GNIH* | Gonadotropin-inhibitory hormone |
| *GNRH1* ** | Gonadotropin releasing hormone 1 |
| *GNRHR2* * | Gonadotropin-releasing hormone receptor |
| *MC2R* | Melanocortin receptor 2 |
| *MC4R* ** | Melanocortin receptor 4 |
| *NR3C1* * | Glucocorticoid receptor |
| *NR3C2* * | Mineralcorticoid receptor |
| *OXTR* * | Oxytocin receptor |
| *POMC* * | Proopiomelanocortin |
| *SERPINA1* | Serpin family A member 1 |
| *VT* | Vasotocin |
| *VTG1* | Vitellogenin-1 |

### Hybridization capture

A pool of 48 libraries contained 39 starling samples, two house sparrow (*Passer domesticus*) samples (to compare bait performance in any conserved regions because we planned to use this bait supplier and protocol with another study on house sparrows), and seven replicate libraries. Eight superb starling samples were used in a previously unpublished DNA methylation study that used RRBS. Three of these eight were included in duplicate libraries and one in triplicate libraries, and the house sparrow samples were also run in duplicate. The remaining 31 additional superb starling samples were included to assess target capture performance when scaling up to a pool of 48 libraries.

We followed the myBaits v4.01 protocol, but lowered the default hybridization and clean up temperature from 65°C to 63°C as myBaits recommends for fragmented DNA. We note, however, that a temperature of 65°C works similarly well with EM-seq converted libraries [24]. Hybridization beads bound to the pooled library-blocker mix were cleaned with three washes, and the resuspended bead-bound DNA was then amplified with 16 PCR cycles in a KAPA HiFi reaction using a 60°C annealing temperature and a one minute extension step at 72°C. The amplified capture pool was subsequently cleaned up with AMPure XP beads and sequenced at 2×150 bp reads with 5% PhiX spike-in in one lane (110 Gb) of an Illumina HiSeq 4000 at the Novogene Corporation, Sacramento, CA.

### Control sequences

Bait library sequence alignments to NEB pUC19 methylated and lambda unmethylated controls were sparse in background coverage because controls were not specifically targeted in our initial bait set. Overall, coverage in control regions ranged from 0-42x without deduplication, averaging 18x for pUC19 and 25x for lambda, but methylation calls were unreliable after deduplication. Although our three EM-seq libraries that were also used for WGEM-Seq passed control checks (mean lambda conversion rate was 1.23%, and mean pUC19 conversion rate was 97.7%), and the RRBS libraries used to compare CpG methylation percentages were spiked with lambda controls, we confirmed that in the seven TEEM-Seq libraries used for comparison to other methods, methylated Cs in CHG and CHH context were very low (0.57% and 0.54%, respectively, on average in alignment reports from Bismark v0.19.0 [25]) as expected, and without the characteristic rise indicating incomplete conversion (typically this rise in CHG/CHH context is between 5 to 10% as in the excluded sample discussed below). Likewise, a known low methylation (average 1.5%) 57 bp region with 12 CpG sites 10 bp upstream of Arginine vasopressin receptor 1A also had means within 1% of this average in the seven TEEM-Seq libraries used for comparison to other methods. Nonetheless, we decided to specifically target pUC19 and lambda control sequences in a parallel TEEM-Seq study using a house sparrow-specific bait set to ensure that we would have reliable EM-seq conversion estimates for all samples in the absence of comparable libraries for these samples prepared with alternative methods [24]. We found that lambda unmethylated and pUC19 methylated control baits were highly specific, targeting the first 1 kb for pUC19 and the first 2 kb for lambda included in the bait design. To ensure requisite control alignments with an untested protocol, and because we only targeted a portion of each control genome, we followed EM-seq recommendations for a higher spike-in with 0.3 ng of pUC19 and 6 ng of lambda controls per 100-150 ng sample. However, given the excessive depth of capture, a spike-in at normal HiSeq levels (a 1:100 dilution, or 0.001 ng pUC19 and 0.02 ng lambda) should be more than sufficient when control regions are targeted. For a capture pool of 96 libraries, SAMtools v1.14 [26] reported coverage for the 1 kb pUC19 alignments as a range of 18,582-538,322x (mean = 96,689x) for non-deduplicated BAMs and a range of 4,540-86,441x (mean = 27,929x) for deduplicated BAMs. For the 2 kb Lambda alignments, SAMtools reported a range of 23,445-537,148x (mean = 115,380x) for non-deduplicated BAMs and a range of 4,984-88,019x (mean = 30,943x) for deduplicated BAMs. Since the bait design and hybridization reaction had no negative effects on the assessment of control alignments, we recommend that all capture designs for TEEM-Seq libraries follow this protocol and include baits for pUC19 and lambda controls.

### TEEM-Seq comparison and validation

To compare and validate TEEM-Seq against more traditional methods of analysis for DNA methylation, we prepared TEEM-Seq libraries from many of the same starling DNA samples that underwent WGEM-Seq and RRBS. For WGEM-Seq, extra material from three of the EM-seq libraries used for bait capture were confirmed clear of adapter dimers on the TapeStation, pooled and sequenced at 2×150 bp reads with 5% PhiX spike-in in 3/4^th^ of a lane (82.5 Gb) on a HiSeq 4000 at Novogene with the same parameters to provide a whole-genome sequence and partial methylome analysis comparison to our targeted sequencing. For a previously unpublished RRBS study, we used NuGEN (now Tecan) Ovation Methyl-Seq kits to prepare RRBS libraries (with added Promega unmethylated cl857 Sam7 lambda control spike-in of 1 ng per 80-120 ng of sample), which were sequenced in several pools of 16 samples per lane (25 Gb) at 1×100 bp reads with 10% PhiX spike-in on a HiSeq 2500 at the Cornell Institute of Biotechnology Genomics Core, Ithaca, NY. Libraries for RRBS were prepared according to kit protocols. Briefly, samples were digested with MspI at 37°C, ligated with bisulfite-compatible barcoded adaptors at 25°C, end repaired at 60°C, bisulfite converted with the Qiagen EpiTect Fast Bisulfite Conversion Kit, PCR amplified with NuGEN’s standard 12 cycles and 60°C annealing, and purified with Agencourt beads for sequencing. Eight of these samples used previously for RRBS were included in the present study, though one failed enzymatic conversion (mean lambda conversion rate from the original RRBS analysis of these samples was 1.43%). Data from the remaining seven samples (from different sequencing pools) were used as a comparison to the TEEM-Seq and WGEM-Seq derived data generated from the same individual starlings.

### DNA methylation analysis

Sequencing data were trimmed with the Trim Galore v0.4.2 [27] wrapper of Cutadapt v1.12 [28] with standard parameters. As per NuGEN’s recommendations, RRBS reads were not trimmed with Trim Galore’s RRBS option, but instead trimmed of adaptors with standard parameters and with NuGEN’s trimRRBSdiversityAdaptCustomers.py script. Trimmed reads were aligned to the superb starling bisulfite genome reference generated by Bismark. WGEM-Seq and TEEM-Seq alignments were deduplicated, and CpG coverage files with methylation percentages (100 * methylated cytosines/total of methylated plus unmethylated cytosines in CpG context) were extracted from alignments with Bismark (non-deduplicated methylation calls were also made for TEEM-Seq data to compare percentages with and without deduplication). Although there does not appear to be a consensus on whether target-enriched libraries should be deduplicated (for amplicon sequencing it is avoided), and because (i) the EM-seq ligation adaptor is specifically designed for the EM-seq prep and (ii) EM-seq currently does not support NEB Unique Molecular Identifiers (which would have to be methylated for compatibility with methylation sequencing workflows), our goal for bait coverage was a minimum of 20x per 4 kb putative promoter target region and 5x for smaller promoter and exonic regions after deduplication.

To compare DNA methylation quantification methods in superb starlings, Bismark coverage files from TEEM-Seq and WGEM-Seq alignments were intersected with 4kb bait “promoter” and exon ranges (for a separate exon comparison) using Bedtools v2.29.2 [29]. To confirm read and CpG site coverage, BAM alignments and Bismark coverage tracks were visually inspected with Geneious v10.0.4 using an updated chromosomal-level genome assembly (CU_Lasu_v2; GenBank: GCA_015883425.2) that combined Hi-C data with the previously published superb starling reference genome [22]. SAMtools coverage was used to calculate mean read coverage stats in bait target ranges. To compare WGEM-Seq and RRBS data, Bedtools was used to generate 200 random 10,000 bp tracks across the starling genome that were then intersected with putative 2.5 kb promoter tracks, leaving us with 27 random “promoter” regions. This targeted 2.5kb range was chosen because RRBS coverage is dense at restriction sites (e.g., in CG clusters localized just prior to the transcription start site). Visual inspection confirmed that these random regions appeared rich in RRBS coverage. Bismark coverage files from WGEM-Seq and RRBS alignments were likewise intersected with these random regions, and then filtered for unique sites (in case of promoter overlap) above 5x coverage [30]. Intersected coverage files for each sample were then matched between each method for common CpG sites. That is, we compared matched sites on a per sample basis, either within random regions for WGEM-Seq versus RRBS data (resulting in 20 regions with 5x CpG site coverage for all but one sample), or within target regions for WGEM-Seq versus RRBS or WGEM-Seq versus TEEM-Seq data. Thus, WGEM-Seq, TEEM-Seq, and RRBS data were compared at the CpG site-level in target regions, and WGEM-Seq and RRBS data were compared at the CpG site-level in 20 random “promoter” regions that were not included in the bait set. Since validation versus older bisulfite methods was done by New England Biolabs themselves in enzymatic conversion development [20], the primary goal of this study was to evaluate TEEM-Seq performance relative to WGEM-Seq and RRBS, which we chose to do using the approach of an additional random 2.5 kb “promoter” region comparison for two reasons. First, we expected that some RRBS or WGEM-Seq reads corresponding to target ranges might have sparse coverage in comparison to the high coverage bait alignments. Second, we wanted to ensure a valid parallel to target regions to check consistency between WGEM-Seq and RRBS outside of target regions, which were also assessed. We chose this approach rather than focusing on sparser areas of common CpG sites because we thought the latter would not necessarily be a good parallel to (namely putative promoter) target regions. Finally, Bedtools map was used to calculate the mean of Bismark methylation percentages for these regions. All alignments were visualized using the Integrative Genomics Viewer (IGV) v2.12.3 [31].

### Statistical analysis

DNA methylation values in target regions for libraries run in duplicate (or triplicate) were compared using paired t-tests. Pearson correlations were used to compare site-level CpG data for TEEM-Seq, WGEM-Seq, and RRBS libraries with the *getCorrelation* function in methylKit v1.20.0 [32]. Data for each pairwise comparison of methods for quantifying DNA methylation (TEEM-Seq versus WGEM-Seq, TEEM-Seq versus RRBS, and WGEM-Seq versus RRBS) were analyzed using generalized linearized mixed models (GLMMS), with DNA methylation as the dependent variable, quantification method as the independent variable, and bird ID included as a random effect to account for repeated measures (multiple gene regions per library). All statistical tests were done in JMP Pro v16.2 or R v4.1.3 [33].

## Results

### Sequencing metrics

To compare sequencing depth investment across methods, we produced summary tables with key metrics (S3-S5 Tables). For TEEM-Seq, our results from a pool of 48 libraries prepared using avian DNA from whole blood in a single capture reaction sequenced on an Illumina HiSeq 4000 at 110 Gb (~2.3 Gb per sample) indicated that target read coverage for the superb starling samples prior to deduplication ranged from a mean of 400-4300x, with an overall mean of 1700x for both DNA strands. Moreover, background, non-target coverage consisted of mostly scattered but evenly distributed single strand alignments, with a mean of just 22x. We determined that while duplication of total reads was up to 95-97% in target regions according to Bismark deduplication reports (versus 15-17% duplicated reads across the genome for the three WGEM-Seq starling libraries, which suggests that sequencing optimization for this protocol might first focus post-capture before reducing pre-capture PCR cycles and per-sample template for the capture reaction), TEEM-Seq target region read coverage depth after deduplication ranged from a mean per target of 20-270x, with an overall mean coverage depth for all targets of 173x (Fig 2). In contrast, WGEM-Seq target region coverage after deduplication ranged from a mean per target of 2-41x, with an overall mean coverage of 15x (Figs 2 and S1 and S3 Tables). In addition, the mean depth of TEEM-Seq coverage after deduplication for just the seven starling samples also included in the RRBS study ranged from 30-374x for the lowest to highest depth putative promoter targets, with an overall mean of 248x (S4 Table). For those same samples’ RRBS libraries, the mean coverage depth per target promoter region (non-deduplication) ranged from 0-17x, with an overall mean of 7x, and a mean CpG coverage of 13.94x across putative promoters (S5 Table). Although RRBS coverage is biased toward restriction sites (C/CGG for MspI) and certain targets had zero coverage, on average 120 CpGs were covered above 5x across strictly 4 kb promoter regions (S6 Table). With the TEEM-Seq and WGEM-Seq libraries, the mean was also coincidentally 120 CpGs above 5x in the three samples compared across methods, though the minimum number of 5x sites covered was never zero for either method (S7 Table). All of the intended target regions were covered in both the TEEM-Seq and WGEM-Seq data, but one target (*DNMT3A;*) was not covered in the RRBS data. Finally, for four samples run as duplicate or triplicate libraries, there were no significant differences in mean DNA methylation between the replicates (all paired t_23_ < 1.63, all P > 0.12; S8 Table), and individual CpG sites were highly correlated within each duplicate or triplicate library (all R = 0.94 to 0.99; S2 Fig).

**Fig 2.**
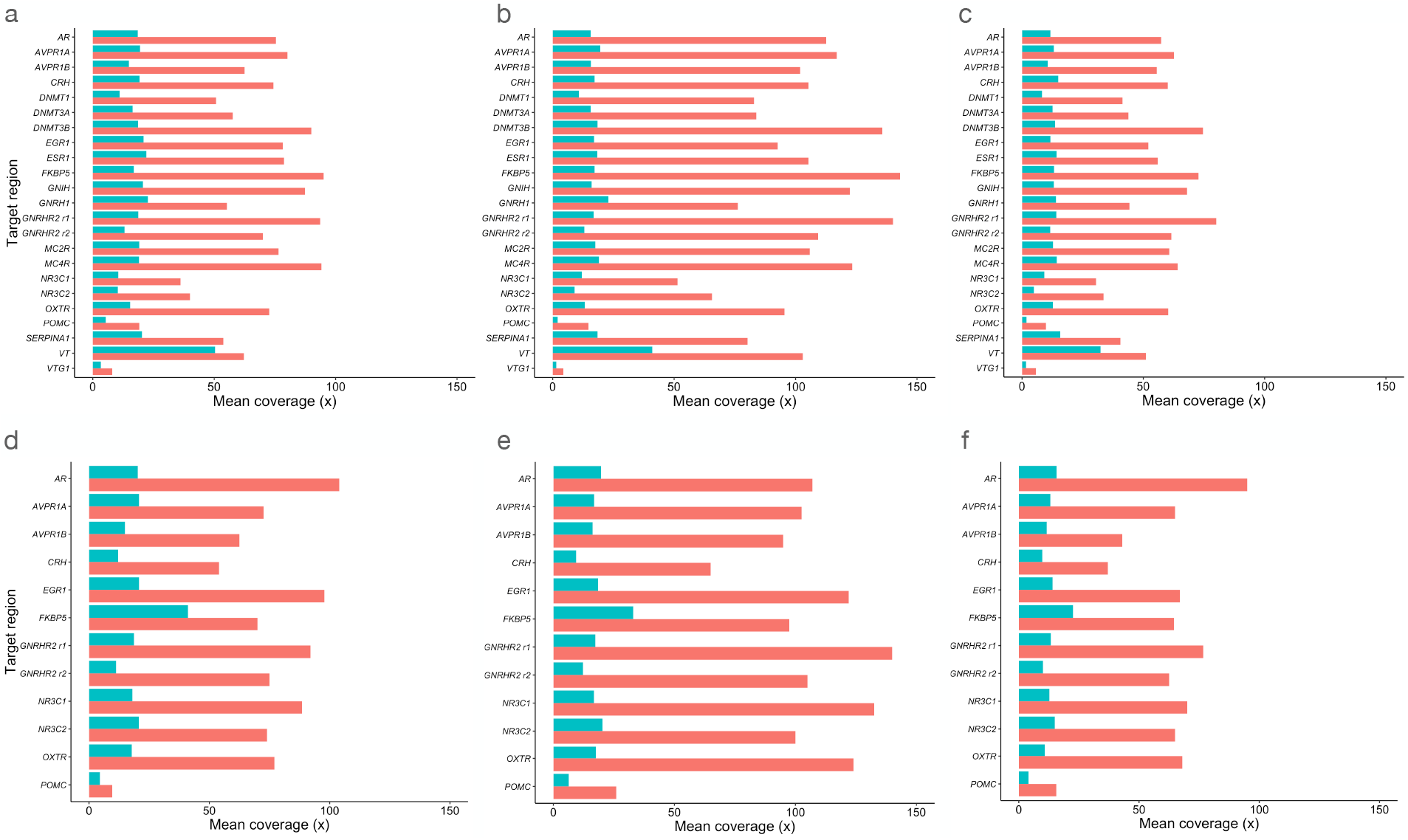
Comparison of mean target region coverage (after deduplication) for target-enriched enzymatic methyl sequencing (TEEM-Seq) and whole-genome enzymatic methyl sequencing (WGEM-Seq) in (a-c) putative promoter and (d-f) exonic regions. Libraries for three of the same individual superb starlings (**a** and **d**) BB-17168, (**b** and **e**) BB-17501, and (**c** and **f**) BB-14232 were sequenced using TEEM-Seq (red bars) and WGEM-Seq (blue bars).

### Assessing bait performance in another avian species

Although our bait set was designed specifically for the superb starling, we included two house sparrow samples (in duplicate) to assess starling-derived bait performance of any conserved regions in another related passerine bird species, and because we intended to use the same bait supplier in a study on house sparrows [24]. For the four house sparrow libraries, SAMtools reported that 19 of 23 putative promoter targets had a mean percent of covered bases of <50%, and 11 of 23 targets had <25% coverage. In contrast, 20 of 23 targets had a mean percent of covered bases >99.5% for the 44 superb starling libraries, with *POMC* having 96% covered bases, and *NR3C1* and *VTG* having reference gaps of 488 bp and 182 bp, respectively, but still >99% coverage elsewhere in the putative promoters. The maximum mean for the target with the highest coverage was 71% in the house sparrow, and the mean depth of coverage after deduplication ranged from a mean across samples of 0-39.5x (versus 20-270x for the superb starling) for the lowest to highest depth putative promoter targets, with an overall mean of 9.3x (versus 173x for the superb starling). As expected, the superb starling bait sequence was not well conserved for most targets in the house sparrow, confirming that species-specific baits should be designed for each study organism using its own reference genome.

### Comparing and validating TEEM-Seq against other methods of DNA methylation analysis

To validate TEEM-Seq against more traditional methods of analysis for DNA methylation, we compared many of the same starling DNA samples prepared for WGEM-Seq and RRBS. At the CpG site-level with a minimum of 5x coverage, TEEM-Seq and WGEM-Seq libraries from the same individual were highly correlated (Fig 3), as were TEEM-Seq and RRBS libraries (Fig 4). Interestingly, when we compared WGEM-Seq and RRBS libraries at a threshold of 5x coverage for CpG sites, the local regression line was a poor fit to the comparison in a number of the samples (S5a Fig). The fit improved when site coverage was increased to 7x (S5b Fig), suggesting that WGEM-Seq and RRBS comparisons required more stringent filtering when looking across the whole genome than did comparisons between WGEM-Seq and TEEM-Seq. In addition, these results imply that TEEM-Seq may be more reliable than RRBS at lower coverage depths, though this should be confirmed by using enzymatic rather than bisulfite conversion for reduced-representation sequencing, something that was beyond the scope of this study.

**Fig 3.**
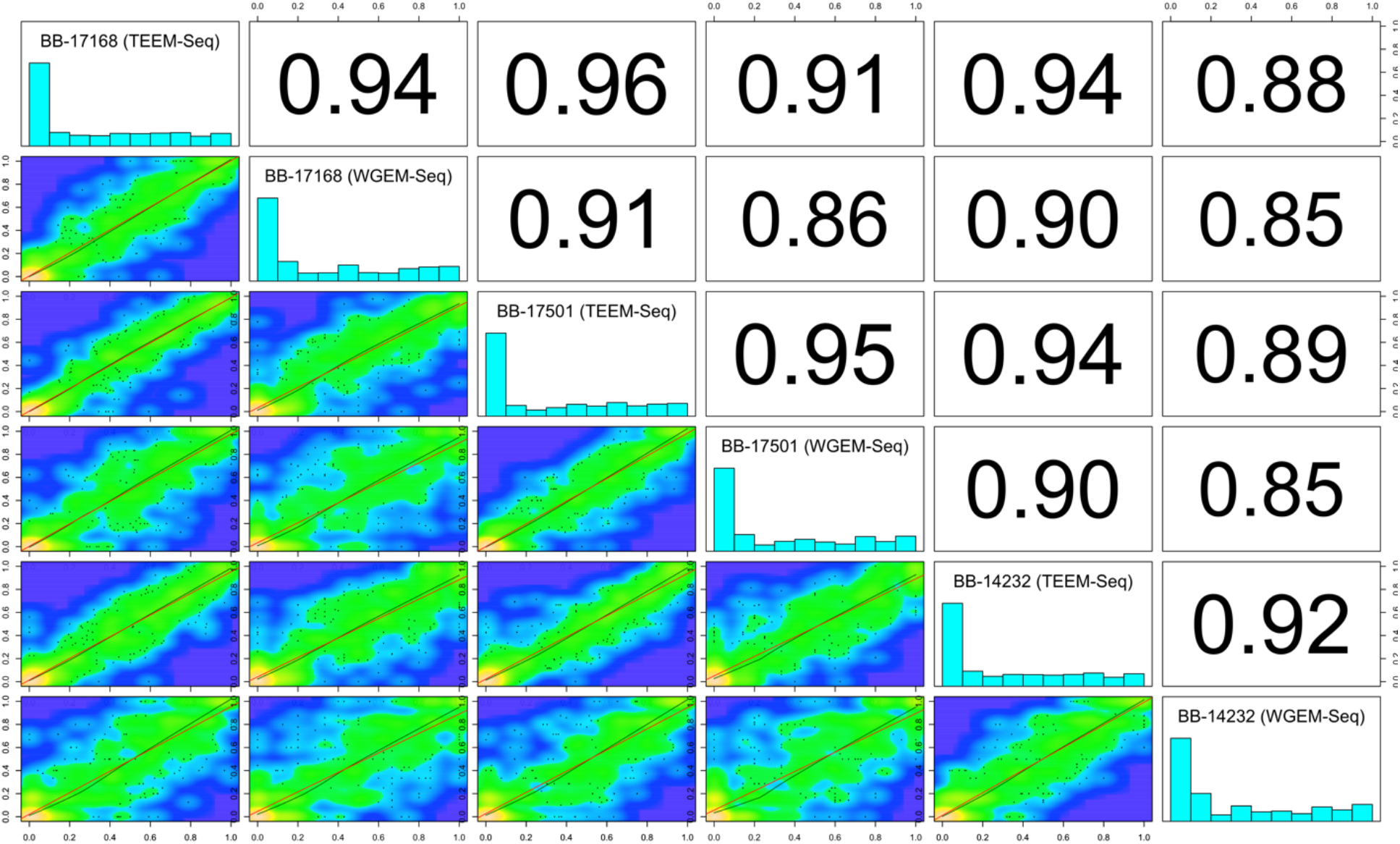
Putative promoter target region CpG site-level percent DNA methylation distribution histograms for each sample library and scatterplots with Pearson correlation coefficients between target-enriched enzymatic methyl sequencing (TEEM-Seq) and wholegenome enzymatic methyl sequencing (WGEM-Seq) libraries from the same individual superb starlings. CpG sites shared between TEEM-Seq and WGEM-Seq libraries at 5x coverage or above were used for comparisons across all libraries (816 CpG sites shared) with the *getCorrelation* function in methylKit for superb starling individuals BB-17168, BB-17501, and BB-14232. Note that S4 Fig shows the same within-individual comparisons, only with higher numbers of shared CpG sites because of within-sample comparisons only. Blue regions of the scatterplots are uncorrelated, yellow regions are highly correlated, and green regions are variably correlated. The green lines represents lowess polynomial regression fits, whereas the red lines represent linear regression fits. Across samples, the correlation coefficients for TEEM-Seq versus TEEM-Seq comparisons are slightly higher (0.95 mean) than for TEEM-Seq versus WGEM-Seq comparisons (0.94 mean), which is not surprising because WGEM-Seq libraries are much lower in mean target region coverage and therefore site-level CpG percent methylation scores could be less reliable; in any case the similarity in correlation coefficient means suggests that TEEM-Seq does not flatten differences in methylation calls as compared with WGEM-Seq. As expected within each sample, the TEEM-Seq versus WGEM-Seq library correlation coefficients are higher for each sample comparison than for WGEM-Seq versus WGEM-Seq compared across samples (see also S6 Fig for WGEM-Seq versus WGEM-Seq complete coverage).

**Fig 4.**
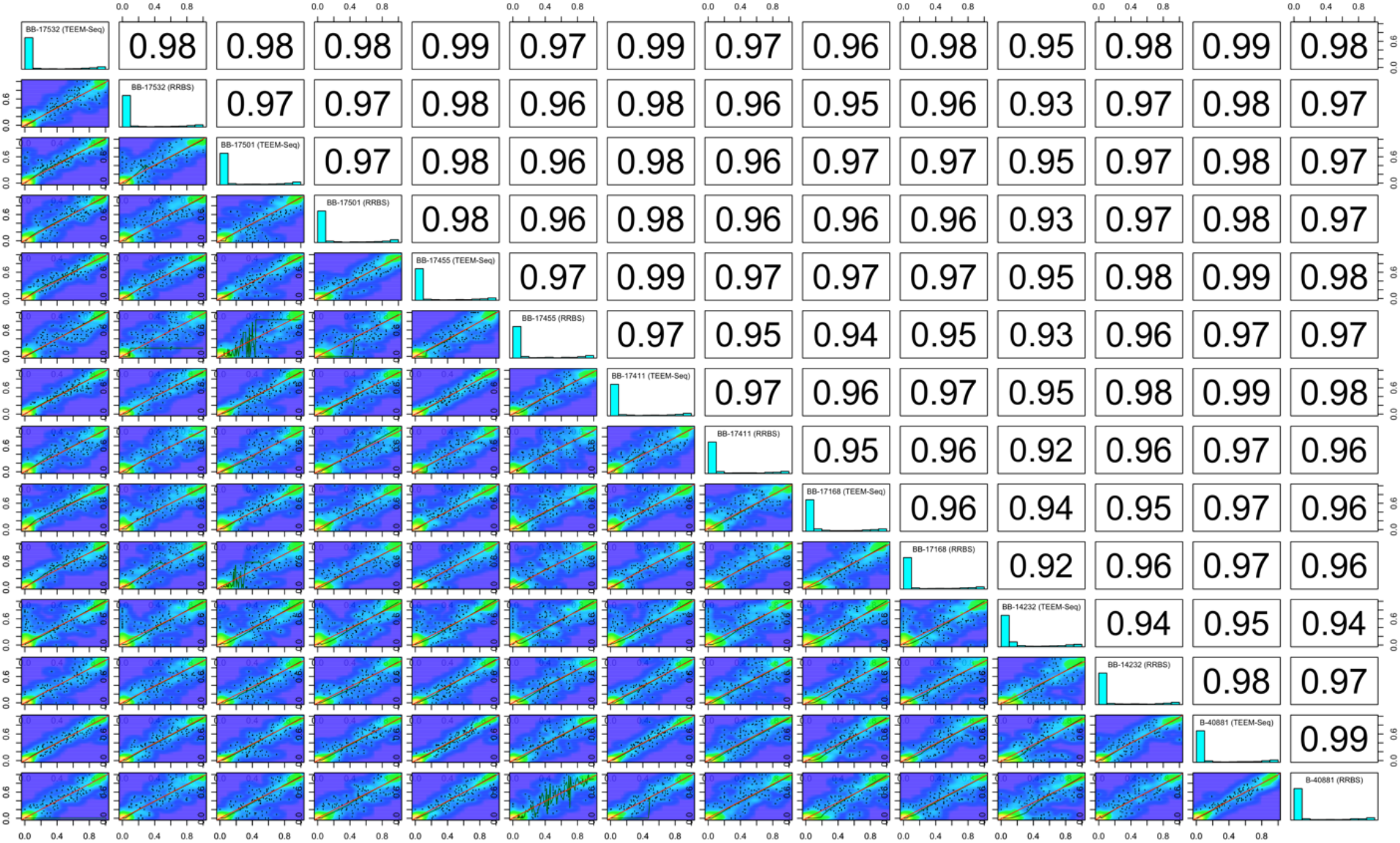
Putative promoter target region CpG site-level percent DNA methylation distribution histograms for each sample library and scatterplots with Pearson correlation coefficients between target-enriched enzymatic methyl sequencing (TEEM-Seq) and reduced-representation bisulfite sequencing (RRBS) libraries from the same individual superb starlings. CpG sites shared between TEEM-Seq and RRBS libraries at 5x coverage or above (1,390 CpG sites shared across all seven samples) were used for paired comparisons with the *getCorrelation* function in methylKit for superb starling individuals BB-17532, BB-17501, BB-17455, BB-17411, BB-17168, BB-14232, and B-40881. Blue regions of the scatterplots are uncorrelated, yellow regions are highly correlated, and green regions are variably correlated. The green lines represents lowess polynomial regression fits, whereas the red lines represent linear regression fits.

We found no major differences in mean DNA methylation across approaches. CpG methylation varied little at the site-level when absolute value mean differences in DNA methylation for sites were compared across gene bodies and putative promoter regions (Figs 5a-c), or when sites were averaged across target regions, as methylation did not differ between WGEM-Seq and TEEM-Seq data within putative promoter (F_1,134_ = 0.0013, P = 0.97; Fig 5d and S7 Table) or exonic regions (F_1,68_ = 0.00001, P = 0.99; S9 Table), between TEEM-Seq and RRBS data within putative promoter regions (F_1,249_ = 0.049, P = 0.83; Fig 5e and S6 Table), or between WGEM-Seq and RRBS in 20 random 2.5 kb “promoter” regions (F_1,114_ = 1.36, P = 0.25; Fig 5f and S10 Table).

**Fig 5.**
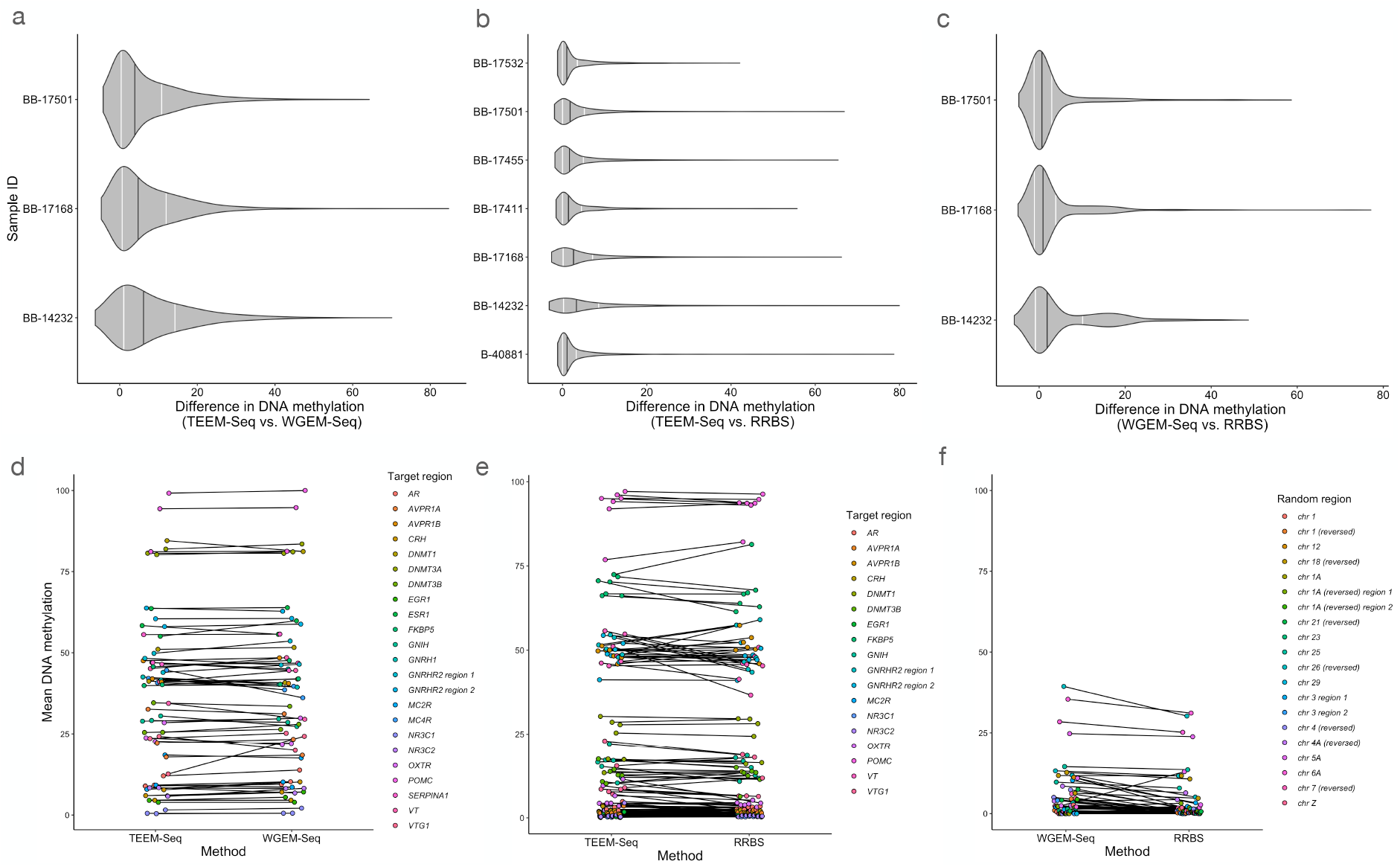
Pairwise comparisons of three methods for quantifying DNA methylation: target-enriched enzymatic methyl sequencing (TEEM-Seq), whole-genome enzymatic methyl sequencing (WGEM-Seq), and reduced-representation bisulfite sequencing (RRBS). (**a** and **d**) Libraries for three of the same individual superb starlings were sequenced using both TEEM-Seq and WGEM-Seq. (**b** and **e**) Libraries for seven of the same individual superb starlings were sequenced using both TEEM-Seq and RRBS. (**c** and **f**) Libraries for three of the same individual superb starlings were sequenced using both WGEM-Seq and RRBS. WGEM-Seq and RRBS data were compared in 20 random 2.5 kb “promoter” regions that were not included in the bait set because we expected that some RRBS and WGEM-Seq reads corresponding to target ranges might have sparse coverage in comparison to the high coverage bait alignments, and as a valid parallel to check consistency between WGEM-Seq and RRBS outside of target regions. In all cases, there were no differences in DNA methylation among quantification methods, as the absolute value mean differences in DNA methylation for promoter regions and gene bodies or random 2.5 kb “promoter” regions did not differ among pairwise comparisons (**a-c**) In each plot, grey violins show the distribution of CpG site absolute value mean differences in DNA methylation for (**a** and **b**) promoter regions and gene bodies or (**c**) random 2.5 kb “promoter” regions, black lines depict the median values, and white lines show the 1^st^ and 3^rd^ quartiles. Additionally, (**d** and **e**) mean DNA methylation levels for each target or (**f**) random region did not significantly differ. Target regions are indicated by different colors according to the legends on panels d-f. See S6-S11 Tables for more detailed information.

Finally, we note that TEEM-Seq data with and without deduplication were nearly identical (S3 Fig) and that despite the high number of duplicates in the TEEM-Seq data, comparisons of WGEM-Seq versus TEEM-Seq were qualitatively similar when using TEEM-Seq data with and without deduplication (S7 and S11 Tables). As expected, CpG site-level correlations were still higher for TEEM-Seq deduplicated versus non-deduplicated data when using the same sites that were highly correlated between TEEM-Seq and WGEM-Seq data (S3 Fig legend and Fig 3). Overall, mean DNA methylation was comparable in all three methods, indicating that TEEM-Seq is an effective approach for quantifying DNA methylation states in specific regions of the genome.

## Discussion

We introduce a flexible, scalable, and inexpensive method for studying DNA methylation in a large number of candidate genes that combines enzymatic conversion of DNA [20] with a custom-designed hybridization capture bait set, and uses starting input DNA as low as 80 ng (S12 Table). Moreover, since EM-Seq will work with as little as 10 ng [20], lower input volumes are likely to work for TEEM-Seq, which may be critical to some applications in non-model organisms where DNA may be limited or non-replenishable. Unlike commercial methods geared primarily for model organisms, TEEM-Seq can be done in a standard academic laboratory environment for any species. The TEEM-Seq approach described here is able to quantify DNA methylation states similarly well to the more traditional approaches of whole-genome and reduced-representation sequencing, and the downstream bioinformatic analysis is essentially the same for all three approaches. We show that TEEM-Seq is both reliable and repeatable, as duplicate libraries from the same samples were highly correlated. Although we demonstrate its effectiveness on DNA from birds, TEEM-Seq should work on DNA from any organism, as well as from DNA extracted from tissues other than blood [24]. And because it employs enzymatic rather than chemical conversion of DNA (i.e., bisulfite treatment), we also expect that TEEM-Seq will be useful for studying formalin-fixed paraffin-embedded (FFPE) samples, as well as degraded or ancient DNA, including samples from museum specimens that are likely to already be partially degraded and only damaged further by traditional bisulfite conversion [34].

In addition to its flexibility (i.e., probes can be designed for varying numbers of genomic regions that can occur anywhere in the genome), one of the greatest strengths of TEEM-Seq is its scalability. Although we only scaled our capture reaction to 48 libraries, we note that in a parallel TEEM-Seq study that applied the method described here using a similar bait set in house sparrows, capture scaled consistently with a 96-library pool [24]. Moreover, New England Biolabs currently produces 120 total unique indexes across their 24- and 96-sample EM-seq kits, meaning that TEEM-Seq should be readily able to scale to a 120-library pool, and perhaps even further with the development of additional unique indexes, or the application of indexes from other vendors. We caution, however, that there is likely to be a tradeoff in the number of baits used in the bait set and the number of samples used in the hybridization capture, though assessing this was beyond the scope of the current study. Since myBaits 80 nt baits are designed to tolerate >20% target divergence in conventional DNA sequencing, it is likely that the eight methylation-state targets myBaits recommended for our bait design, in addition to the unmodified targets, were to some degree redundant. If this proves to be the case, costs could be reduced further in bait design, though assessing the capture performance and mismatch allowance of the unmodified baits and redundancy of the methylation-state bait permutations was beyond the scope of this study. Although we cannot directly assess bias in whether and how specific baits hybridize and whether this could have an effect on methylation levels, under the current design parameters, no systemic bias was evident in capture products. Ultimately, TEEM-Seq is highly scalable because the conversion of unmethylated cytosines to uracils is done prior to the hybridization reaction. In contrast, Agilent’s SureSelect^XT^ Methyl-Seq Target Enrichment System performs hybridization capture before library pooling, which limits the number of samples to a fraction of that possible in each TEEM-Seq capture reaction [17]. While Roche’s SeqCap Epi Enrichment System [18] performs hybridization capture after bisulfite conversion, it is also currently limited to a fraction of those indexes available for EM-seq, which similarly constrains the number of samples in each capture reaction. Finally, we note that Twist Bioscience has recently developed custom methylation probe panels for target enrichment that also works with EM-Seq, though this approach has thus far only been optimized for humans, and to our knowledge, there are currently no published studies validating this method in other species (https://www.twistbioscience.com/resources/technical-note/highly-sensitive-methylation-detection-using-enzymatic-methyl-seq-and).

Finally, TEEM-Seq is more cost-effective and per sample data-efficient than not only other commercial capture-based approaches, but also whole-genome sequencing (S12 Table). TEEM-Seq is also comparable in cost to reduced-representation sequencing; TEEM-Seq library prep was less than one third the cost (roughly $25 per sample, including probes, sample quality control, and costs associated with DNA shearing) of our RRBS study, which used NuGEN (Tecan) Ovation Methyl-Seq kits. For an avian species with a genome size of roughly 1.07 Gb [22], we sequenced approximately 2.5 Gb per sample for TEEM-Seq, 1.75 Gb per sample for RRBS, and 27.5 Gb per sample for WGEM-Seq. We note, however, that even with an order of magnitude lower sequencing demand for TEEM-Seq compared to WGEM-Seq, the specificity of the bait set meant that we routinely produced 100x coverage or more of the targets, including control regions, and all target regions had mean coverage of at least 20x after deduplication. Given how we suspect from the correlation results that TEEM-Seq performed better than RRBS at 5x coverage compared to WGEM-Seq, we may have actually needed slightly greater depth in RRBS than we sequenced. Sequencing depth for TEEM-Seq could be further reduced with additional optimization and perhaps with a reduction in bait redundancy. In addition, since we were able to reduce EM-seq kit reagents down to three quarter volumes and have similar success, by maximizing the number of samples per total capture pool (up to 120 samples per reaction with current index availability), TEEM-Seq becomes an inexpensive method for accurately quantifying DNA methylation states across continuous regions of the genome in a large sample for a diversity of organisms. Finally, all three methods are roughly similar in terms of library preparation time (assuming 96 samples), with RRBS and WGEM-Seq taking between three and four days and TEEM-Seq five days because of the added capture step (S12 Table). For the same number of reads, TEEM-Seq results in higher coverage for target regions than WGEM-Seq. Whereas for RRBS, regions of interest will be missed (*DNMT3A* in our gene set had to be excluded from RRBS comparisons due to very sparse coverage) because of inherent bias toward CpG islands that may or may not fall within promoters for a given pathway. In addition RRBS will only sample a few CpG clusters in regions of interest, whereas TEEM-Seq can be designed to samples all of the CpG sites within those regions.

We expect that TEEM-Seq could replace traditional approaches for studying candidate genes and pathways (e.g., amplicon sequencing or pyrosequencing), which are time and labor intensive, and be effectively paired with other DNA methylation quantification methods (e.g., whole-genome or reduced-representation sequencing) to increase project sample sizes. Moreover, TEEM-Seq can be combined with mRNA sequencing to examine how DNA methylation in promoters or other regulatory regions is related to the expression of individual genes or gene networks. As EM-seq and other enzymatic approaches for conversion of DNA continue to expand and replace bisulfite conversion and sequencing [35–38], we expect that TEEM-Seq can be further optimized to improve efficiency, scale to ever larger numbers of samples, and use other bait set designs. Ultimately, because of its flexibility, scalability, and cost-effectiveness, TEEM-Seq should prove useful to researchers in a number of biological disciplines for rapidly quantifying DNA methylation states in any organism of interest in order to link epigenetics to the development of phenotypes.

## Acknowledgements

We thank W. Watetu, G. Manyaas, and J. Mosiany for assistance collecting superb starlings and the Mpala Research Centre for logistical support in Kenya. We acknowledge A. Devault and Z. Hanf at Daicel Arbor Biosciences for help designing the bait sets. We acknowledge the Sackler Institute for Comparative Genomics at the American Museum of Natural History and the Department of Biological Sciences at Columbia University for access to equipment. We thank S. Siller for helping to organize gene sequences and, along with B. Heidinger, for supplying house sparrow DNA. A. Bendesky and J. Merritt provided constructive feedback on previous versions of this manuscript.

## Data Availability

Raw genetic data are available at the NCBI Sequence Read Archive (SRA) (BioProject: PRJNA861281). Summarized data are provided in the Supplemental Information in table format. Code is available at https://doi.org/10.6084/m9.figshare.20634759.

## Author Contributions

D.R.R. conceived of the project. D.R.R. and J.S. designed the experiment. D.R.R. collected the samples. J.S. collected and analyzed the data. D.R.R. and J.S. wrote the paper.

## References

1. Waddington CH. The epigenotype. Endeavor. 1942; 81:18–20.

2. Jablonka E, Lamb MJ. The changing concept of epigenetics. Ann N Y Acad Sci. 2002; 981:82–96.

3. Cavalli G, Heard E. Advances in epigenetics link genetics to the environment and disease. Nature. 2019; 571:489–499.

4. Rey O, Eizaguirre C, Angers B, Baltazar-Soares M, Sagonas K, Prunier JG et al. Linking epigenetics and biological conservation: Towards a conservation epigenetics perspective. Funct Ecol. 2020; 34:414–427.

5. Robertson KD, Wolffe AP. DNA methylation in health and disease. Nat Rev Genet. 2000; 1:11–19.

6. Bossdorf O, Richards CL, Pigliucci M. Epigenetics for ecologists. Ecol Lett. 2007; 11:106-115.

7. Bird AP. CpG-rich islands and the function of DNA methylation. Nature. 1986; 321:209–213.

8. Harris RA, Wang T, Coarfa C, Nagarajan RP, Hong C, Downey SL et al. Comparison of sequencing-based methods to profile DNA methylation and identification of monoallelic epigenetic modifications. Nat Biotechnol. 2010; 28:1097–1105.

9. Bock C, Tomazou EM, Brinkman AB, Müller F, Simmer F, Gu H et al. Quantitative comparison of genome-wide DNA methylation mapping technologies. Nat Biotechnol. 2010; 28:1106–1114.

10. consortium BLUEPRINT, Bock C, Halbritter F, Carmona FJ, Tierling S, Datlinger P et al. Quantitative comparison of DNA methylation assays for biomarker development and clinical applications. Nat Biotechnol. 2016; 34:726–737.

11. Crary-Dooley FK, Tam ME, Dunaway KW, Hertz-Picciotto I, Schmidt RJ, LaSalle JM. A comparison of existing global DNA methylation assays to low-coverage whole-genome bisulfite sequencing for epidemiological studies. Epigenetics. 2017; 12:206–214.

12. Gouil Q, Keniry A. Latest techniques to study DNA methylation. Essays Biochem. 2019; 63:639–648.

13. Husby A. Wild epigenetics: insights from epigenetic studies on natural populations. Proceedings of the Royal Society B: Biological Sciences. 2022; 289:20211633.

14. Gu H, Smith ZD, Bock C, Boyle P, Gnirke A, Meissner A. Preparation of reduced representation bisulfite sequencing libraries for genome-scale DNA methylation profiling. Nature Protocols. 2011; 6:468–481.

15. Tanaka K, Okamoto A. Degradation of DNA by bisulfite treatment. Bioorg Med Chem Lett. 2007; 17:1912–1915.

16. Kozarewa I, Armisen J, Gardner AF, Slatko BE, Hendrickson CL. Overview of Target Enrichment Strategies. Current Protocols in Molecular Biology. 2015; 112:7.21.1–7.21.23.

17. Lee E-J, Pei L, Srivastava G, Joshi T, Kushwaha G, Choi J-H et al. Targeted bisulfite sequencing by solution hybrid selection and massively parallel sequencing. Nucleic Acids Res. 2011; 39:e127–e127.

18. Li Q, Suzuki M, Wendt J, Patterson N, Eichten SR, Hermanson PJ et al. Post-conversion targeted capture of modified cytosines in mammalian and plant genomes. Nucleic Acids Res. 2015; 43:e81–e81.

19. Masser DR, Stanford DR, Hadad N, Giles CB, Wren JD, Sonntag WE et al. Bisulfite oligonucleotide-capture sequencing for targeted base-and strand-specific absolute 5-methylcytosine quantitation. AGE. 2016; 38:49.

20. Vaisvila R, Ponnaluri VKC, Sun Z, Langhorst BW, Saleh L, Guan S et al. Enzymatic methyl sequencing detects DNA methylation at single-base resolution from picograms of DNA. Genome Res. 2021; 31:1280–1289.

21. Sapolsky RM, Romero LM, Munck AU. How do glucocorticoids influence stress responses? Integrating permissive, suppressive, stimulatory, and preparative actions. Endocr Rev. 2000; 21:55–89.

22. Rubenstein DR, Corvelo A, MacManes MD, Maia R, Narzisi G, Rousaki A et al. Feather gene expression elucidates the developmental basis of plumage iridescence in African starlings. J Hered. 2021; 112:417–429.

23. Seutin G, White BN, Boag PT. Preservation of avian blood and tissue samples for DNA analyses. Can J Zool. 1991; 69:82–90.

24. Siller SJ Epigenetic modification of the hypothalamic-pituitary-adrenal axis during early life of the house sparrow (Passer domesticus). Department of Ecology, Evolution and Environmental Biology 2022; Ph.D.:

25. Krueger F, Andrews SR. Bismark: a flexible aligner and methylation caller for Bisulfite-Seq applications. Bioinformatics. 2011; 27:1571–1572.

26. Danecek P, Bonfield JK, Liddle J, Marshall J, Ohan V, Pollard MO et al. Twelve years of SAMtools and BCFtools. GigaScience. 2021; 10:giab008.

27. Krueger F, James F, Ewels P, Afyounian E, Schuster-Boeckler B. Trim Galore v0.4.2. 2021;

28. Martin M. Cutadapt removes adapter sequences from high-throughput sequencing reads. EMBnetjournal. 2011; 17:10–12.

29. Quinlan AR, Hall IM. BEDTools: a flexible suite of utilities for comparing genomic features. Bioinformatics. 2010; 26:841–842.

30. Ziller MJ, Hansen KD, Meissner A, Aryee MJ. Coverage recommendations for methylation analysis by whole-genome bisulfite sequencing. Nature Methods. 2015; 12:230–232.

31. Robinson JT, Thorvaldsdóttir H, Winckler W, Guttman M, Lander ES, Getz G et al. Integrative genomics viewer. Nat Biotechnol. 2011; 29:24–26.

32. Akalin A, Kormaksson M, Li S, Garrett-Bakelman FE, Figueroa ME, Melnick A et al. methylKit: a comprehensive R package for the analysis of genome-wide DNA methylation profiles. Genome Biol. 2012; 13:R87.

33. Team RC. R: A Language and environment for statistical computing. 2022;

34. Rubi TL, Knowles LL, Dantzer B. Museum epigenomics: Characterizing cytosine methylation in historic museum specimens. Molecular Ecology Resources. 2020; 20:1161–1170.

35. Feng S, Zhong Z, Wang M, Jacobsen SE. Efficient and accurate determination of genome-wide DNA methylation patterns in *Arabidopsis thaliana with enzymatic methyl sequencing*. Epigenetics & Chromatin. 2020; 13:42.

36. Han Y, Zheleznyakova GY, Marincevic-Zuniga Y, Kakhki MP, Raine A, Needhamsen M et al. Comparison of EM-seq and PBAT methylome library methods for low-input DNA. Epigenetics. 2022; 17:1195–1204.

37. Liu Y, Hu Z, Cheng J, Siejka-Zielińska P, Chen J, Inoue M et al. Subtraction-free and bisulfite-free specific sequencing of 5-methylcytosine and its oxidized derivatives at base resolution. Nature Communications. 2021; 12:618.

38. Morrison J, Koeman JM, Johnson BK, Foy KK, Beddows I, Zhou W et al. Evaluation of whole-genome DNA methylation sequencing library preparation protocols. Epigenetics & Chromatin. 2021; 14:28.

