## Supplemental Information for "Target-enriched enzymatic methyl sequencing: flexible, scalable and inexpensive hybridization capture for quantifying DNA methylation"

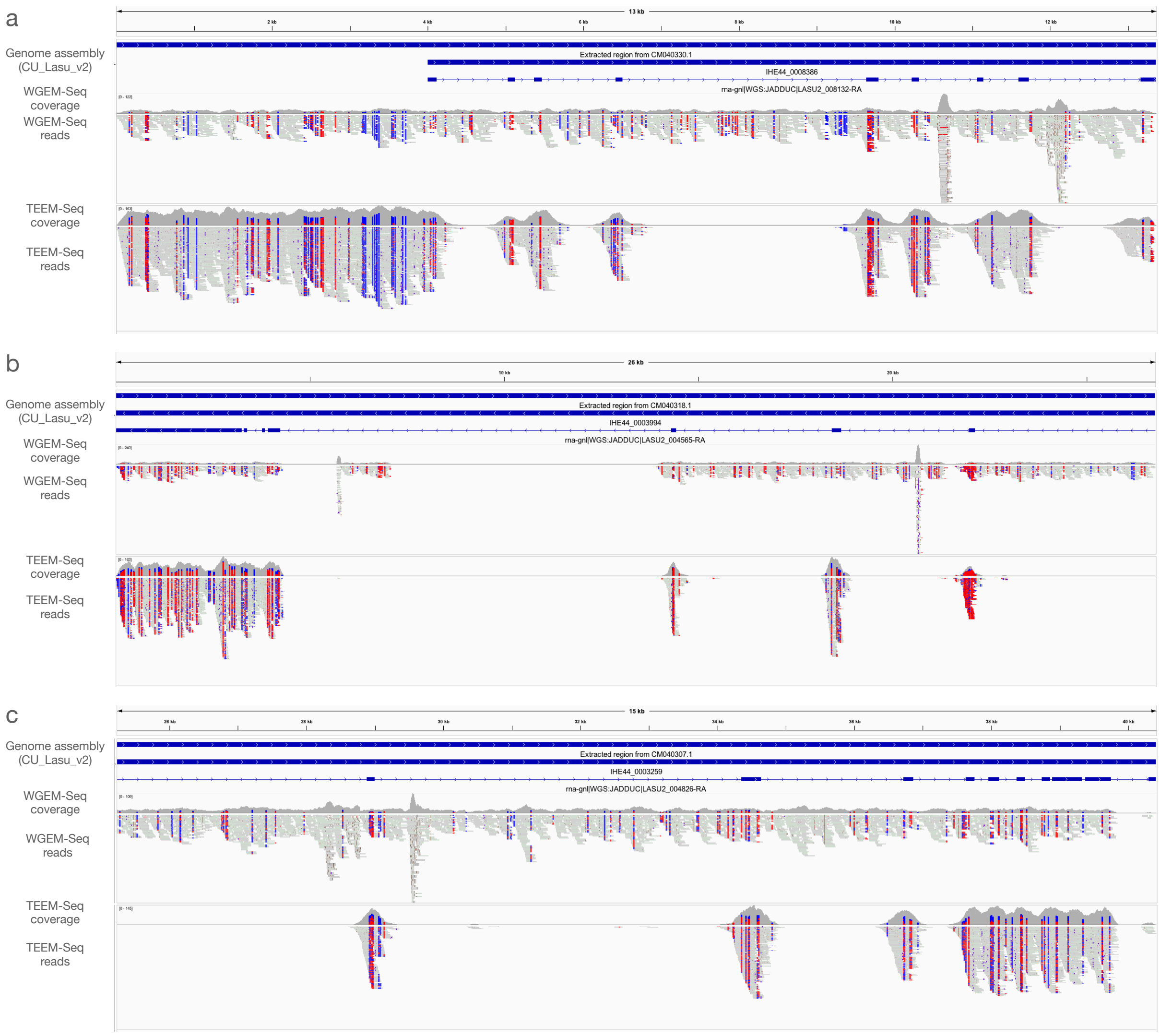


**S1 Fig. Representative examples of target-enriched enzymatic methyl sequencing (TEEM-Seq) versus whole-genome enzymatic methyl sequencing (WGEM-Seq) coverage**. Integrative Genomics Viewer (IGV) logarithm-scale coverage and read alignment plots (for deduplicated data) of superb starling sample BB-17168 multi-exonic targets for (**a**) FK506 binding protein 5 (*FKBP5*), (**b**) the glucocorticoid receptor (*NR3C1*), and (**c**) the androgen receptor (*AR*). *FKBP5* includes the entire promoter and gene region, whereas *NR3C1* and *AR* are focused on the end of each gene region to enable IGV's display of read alignments. Alignments are colored in bisulfite mode by CG, with red indicating methylated sites (non-converted cytosines) and blue indicating unmethylated sites (converted cytosines). Target tracks for the superb starling genome assembly (updated chromosomal-level genome assembly: CU_Lasu_v2; GenBank: GCA_015883425.2) include GenBank sequence ID, locus tag, and protein IDs (with exonic spans also in blue).

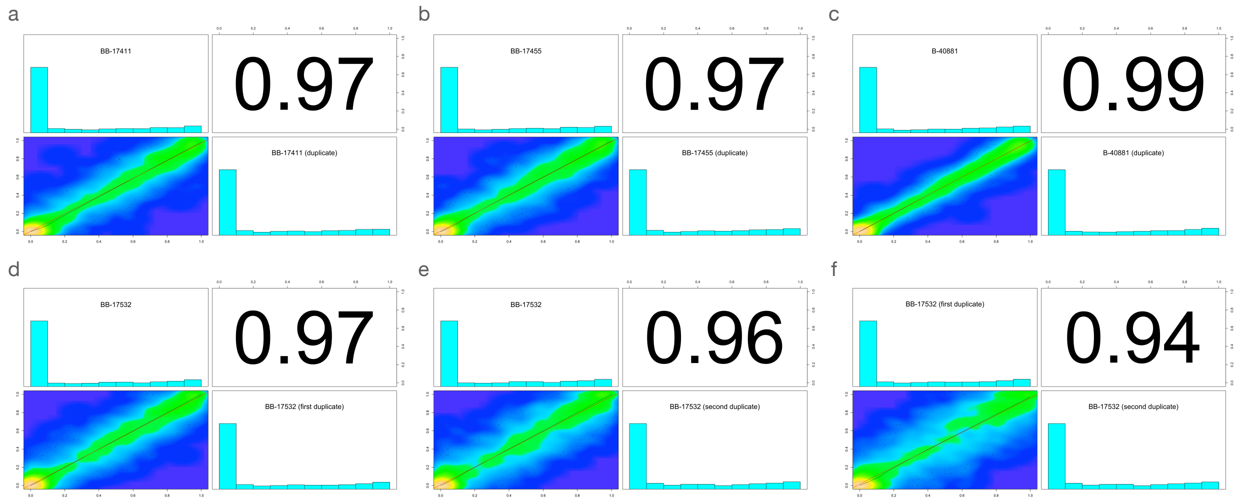


**S2 Fig. Putative promoter target region CpG site-level percent DNA methylation distribution histograms for each sample library and scatterplots with Pearson correlation coefficients between target-enriched enzymatic methyl sequencing (TEEM-Seq) duplicate and triplicate libraries for superb starling individuals BB-17411, BB-17455, B-40881, and BB-17532.** CpG sites shared between TEEM-Seq libraries at 5x coverage or above were used for paired comparisons within samples with the *getCorrelation* function in methylKit: (a) BB-17411 has 5,073 CpG sites shared with its duplicate library; (b) BB-17455 has 5,034 CpG sites shared with its duplicate library; (c) B-40881 has CpG 5,410 sites shared with its duplicate library; and (d) BB-17532 has 4,867 CpG sites shared with its first duplicate library, (e) 4,309 CpG sites shared with its second duplicate library, and (e) 4,248 CpG sites shared between the first and second duplicate libraries. Blue regions of the scatterplots are uncorrelated, yellow regions are highly correlated, and green regions are variably correlated. The green lines represents lowess polynomial regression fits, whereas the red lines represent linear regression fits.

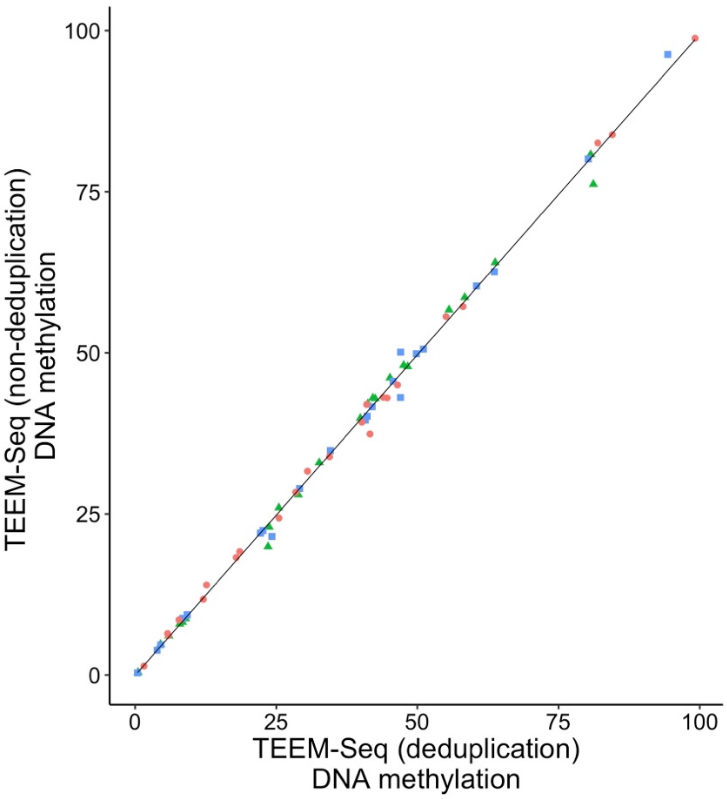


**S3 Fig. Comparison of target-enriched enzymatic methyl sequencing (TEEM-Seq) data with and without deduplication.** Libraries for three individual superb starlings (BB-17168 [green triangles], BB-17501 [blue squares], and BB-14232 [red circles]) were analyzed with and without deduplication. Mean DNA methylation by putative promoter target region for deduplication and non-deduplication of samples were similar (the mean absolute difference across genes was 0.77 for BB-17168, 0.78 for BB-17501, and 0.89 for BB-14232). To parallel Fig 5a, CpG site-level Pearson correlations were also analyzed using the same sites for BB-17168 (R = 0.99, N = 2733, P < 0.0001), BB-17501 (R = 0.99, N = 2770, P < 0.0001), and BB-14232 (R = 0.99, N = 1588, P < 0.0001).

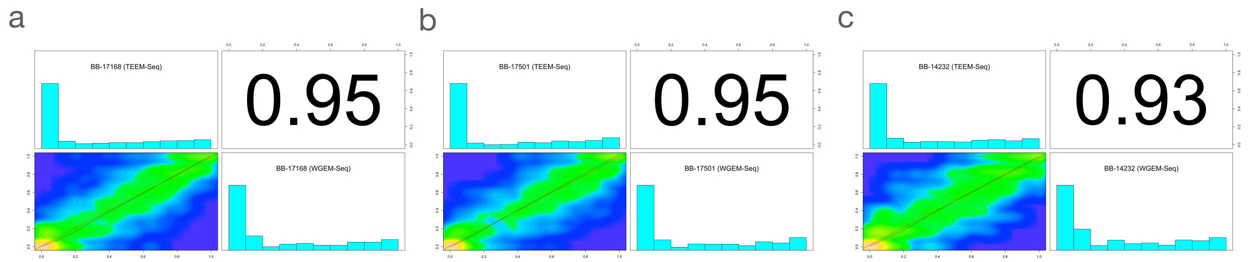


**S4 Fig. Putative promoter target region CpG site-level percent DNA methylation distribution histograms for each sample library and scatterplots with Pearson correlation coefficients between target-enriched enzymatic methyl sequencing (TEEM-Seq) and whole-genome enzymatic methyl sequencing (WGEM-Seq) libraries from the same individual superb starlings.** CpG sites shared between TEEM-Seq and WGEM-Seq libraries at 5x coverage or above were used for paired comparisons within samples with the *getCorrelation* function in methylKit for (a) BB-17168 (2,732 CpG sites shared), (b) BB-17501 (2,769 CpG sites shared), and (c) BB-14232 (1,587 CpG sites shared). Blue regions of the scatterplots are uncorrelated, yellow regions are highly correlated, and green regions are variably correlated. The green lines represents lowess polynomial regression fits, whereas the red lines represent linear regression fits.

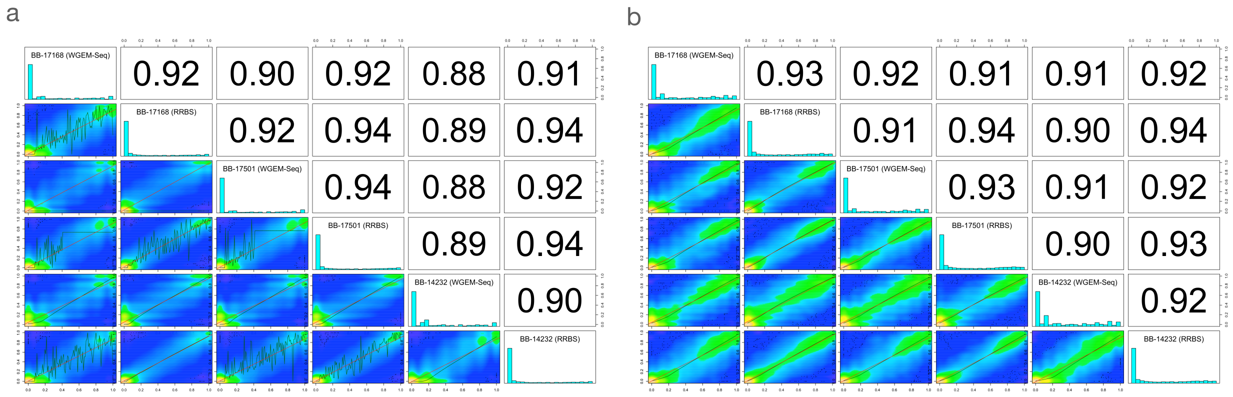


**S5 Fig. Comparisons of CpG site-level percent DNA methylation distribution histograms for each sample library and scatterplots with Pearson correlation coefficients of whole-genome enzymatic methyl sequencing (WGEM-Seq) and reduced-representation bisulfite sequencing (RRBS) libraries from the same three individual superb starlings: BB-17168, BB-17501, and BB-14232.** CpG sites shared at 5x coverage or above were used for paired comparisons with the *getCorrelation* function in methylKit. (a) Comparisons of the entire genome at a 5x threshold for CpG site coverage include 271,828 sites shared across all libraries, whereas (b) comparisons at a 7x threshold include 37,611 CpG sites shared across all libraries. Blue regions of the scatterplots are uncorrelated, yellow regions are highly correlated, and green regions are variably correlated. The green lines represents lowess polynomial regression fits, whereas the red lines represent linear regression fits.

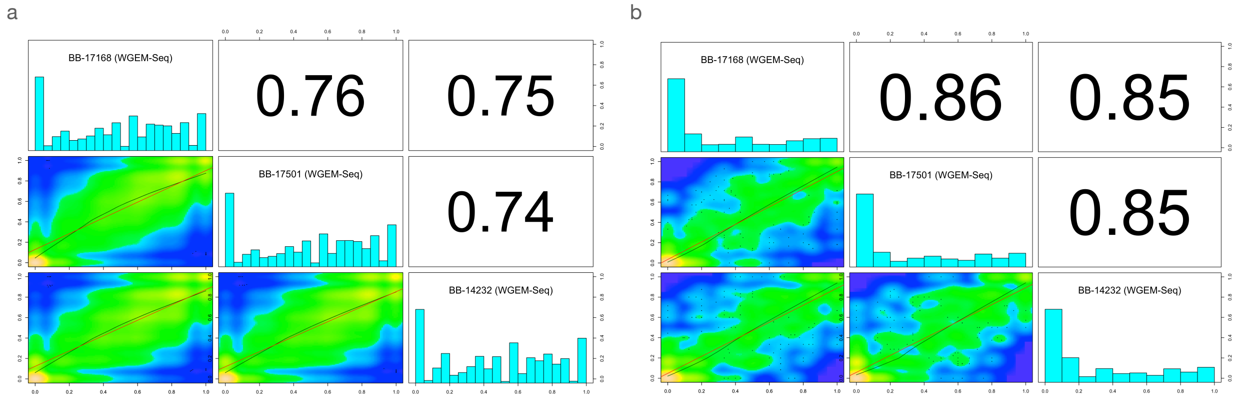


**S6 Fig. Comparisons of CpG site-level percent DNA methylation distribution histograms for each sample library and scatterplots with Pearson correlation coefficients of whole-genome enzymatic methyl sequencing (WGEM-Seq) libraries from the same three individual superb starlings: BB-17168, BB-17501, and BB-14232.** CpG sites shared across WGEM-Seq libraries at 5x coverage or above were used for paired comparisons with the *getCorrelation* function in methylKit. (a) Comparisons of the entire genome include 5,017,620 CpG sites shared across all libraries, whereas (b) comparisons in just the probe target regions include 831 CpG sites shared across all libraries. Blue regions of the scatterplots are uncorrelated, yellow regions are highly correlated, and green regions are variably correlated. The green lines represents lowess polynomial regression fits, whereas the red lines represent linear regression fits.

#### S1 Table. Location of putative promoter bait targets in the superb starling reference genome.

| Target region | Putative promoter region (bp) | Genome position | | Putative promoter bait target | |
|---|---|---|---|---|---|
| | | CU_Lasu_v2 sequence | Chromosome | Start sequence location (1-based) | End sequence location |
| *AR* | 4000 | CM040307.1 | 4A | 14532331 | 14536330 |
| *AVPR1A* | 4000 | CM040303.1 | 1A | 45531591 | 45535590 |
| *AVPR1B* | 4000 | CM040330.1 | 26 | 5619147 | 5623146 |
| *CRH* | 4000 | CM040304.1 | 2 | 116333274 | 116337273 |
| *DNMT1* | 4000 | JADDUC020000037.1 | N/A | 142444 | 146443 |
| *DNMT3A* | 4000 | CM040305.1 | 3 | 111778701 | 111782700 |
| *DNMT3B* | 4000 | CM040324.1 | 20 | 6045685 | 6049684 |
| *EGR1* | 4000 | CM040318.1 | 13 | 19257051 | 19261050 |
| *ESR1* | 4000 | CM040305.1 | 3 | 54837600 | 54841599 |
| *FKBP5* | 4000 | CM040330.1 | 26 | 2992197 | 2996196 |
| *GNIH* | 4000 | CM040304.1 | 2 | 32620135 | 32624134 |
| *GNRH1* | 5726* | CM040326.1 | 22 | 2727612 | 2733338 |
| *GNRHR2 r1*[#] | 4000 | CM040315.1 | 10 | 17836010 | 17840009 |
| *GNRHR2 r2*[#] | 4000 | CM040315.1 | 10 | 20554109 | 20558108 |
| *MC2R* | 3856 | CM040304.1 | 2 | 96963447 | 96967302 |
| *MC4R* | 3948* | CM040304.1 | 2 | 70409831 | 70413779 |
| *NR3C1* | 4000 | CM040318.1 | 13 | 18404677 | 18408676 |
| *NR3C2* | 4000 | CM040306.1 | 4 | 59434518 | 59438517 |
| *OXTR* | 4000 | CM040317.1 | 12 | 20204345 | 20208344 |
| *POMC* | 258** | JADDUC020000174.1 | N/A | 1 | 258 |
| *SERPINA1* | 4000 | CM040308.1 | 5A | 15667895 | 15671894 |
| *VT* | 4000 | CM040306.1 | 4 | 3393601 | 3397600 |
| *VTG1* | 697** | CM040313.1 | 8 | 18775171 | 18775867 |

[*] Since there is no existing annotation in the superb starling reference genome for this gene, we used the zebra finch sequence alignment to target 2 kb upstream in the putative promoter region and 2 kb in the gene body.
[**] Limited sequence upstream of gene.
[#] Two separate gene regions (indicated as r1 and r2) on chromosome 10 with similarity to *GNRHR2*.

#### S2 Table. Location of exon bait targets in the superb starling reference genome.

| Target region | Genome position *CU_Lasu_v2 sequence* | Exon start sequence location (1-based) | Exon end sequence location |
|---|---|---|---|
| *AR* | CM040307.1 | 14536331 | 14536608 |
| *AR* | CM040307.1 | 14553510 | 14553661 |
| *AR* | CM040307.1 | 14561214 | 14561330 |
| *AR* | CM040307.1 | 14566681 | 14566968 |
| *AR* | CM040307.1 | 14569050 | 14569194 |
| *AR* | CM040307.1 | 14569959 | 14570089 |
| *AR* | CM040307.1 | 14570290 | 14570447 |
| *AR* | CM040307.1 | 14570702 | 14570827 |
| *AR* | CM040307.1 | 14571069 | 14571192 |
| *AR* | CM040307.1 | 14571221 | 14571654 |
| *AR* | CM040307.1 | 14571708 | 14572081 |
| *AR* | CM040307.1 | 14572630 | 14572735 |
| *AVPR1A* | CM040303.1 | 45528656 | 45528936 |
| *AVPR1A* | CM040303.1 | 45530634 | 45531541 |
| *AVPR1A* | CM040303.1 | 45531556 | 45531590 |
| *AVPR1B* | CM040330.1 | 5617298 | 5617587 |
| *AVPR1B* | CM040330.1 | 5618209 | 5619146 |
| *CRH* | CM040304.1 | 116332740 | 116333273 |
| *EGR1* | CM040318.1 | 19261051 | 19261285 |
| *EGR1* | CM040318.1 | 19261584 | 19262881 |
| *FKBP5* | CM040330.1 | 2996197 | 2996314 |
| *FKBP5* | CM040330.1 | 2997227 | 2997320 |
| *FKBP5* | CM040330.1 | 2997561 | 2997662 |
| *FKBP5* | CM040330.1 | 2998610 | 2998702 |
| *FKBP5* | CM040330.1 | 3001834 | 3001990 |
| *FKBP5* | CM040330.1 | 3002418 | 3002508 |
| *FKBP5* | CM040330.1 | 3003256 | 3003339 |
| *FKBP5* | CM040330.1 | 3003787 | 3003920 |
| *FKBP5* | CM040330.1 | 3005358 | 3005553 |
| *GNRHR2 r1* | CM040315.1 | 17840010 | 17840015 |
| *GNRHR2 r1* | CM040315.1 | 17840516 | 17841025 |
| *GNRHR2 r1* | CM040315.1 | 17841058 | 17841302 |
| *GNRHR2 r1* | CM040315.1 | 17841288 | 17841489 |
| *GNRHR2 r1* | CM040315.1 | 17841772 | 17842178 |
| *GNRHR2 r1* | CM040315.1 | 17842084 | 17842174 |
| *GNRHR2 r2* | CM040315.1 | 20552342 | 20553124 |
| *GNRHR2 r2* | CM040315.1 | 20553536 | 20554108 |
| *NR3C1* | CM040318.1 | 18337403 | 18340650 |
| *NR3C1* | CM040318.1 | 18340700 | 18340793 |
| *NR3C1* | CM040318.1 | 18341168 | 18341240 |
| *NR3C1* | CM040318.1 | 18341319 | 18341645 |
| *NR3C1* | CM040318.1 | 18351707 | 18351837 |
| *NR3C1* | CM040318.1 | 18355842 | 18356086 |
| *NR3C1* | CM040318.1 | 18359382 | 18359539 |
| *NR3C1* | CM040318.1 | 18365651 | 18365767 |
| *NR3C1* | CM040318.1 | 18369957 | 18370120 |
| *NR3C1* | CM040318.1 | 18403494 | 18404676 |
| *NR3C2* | CM040306.1 | 59422547 | 59422562 |
| *NR3C2* | CM040306.1 | 59430594 | 59432361 |
| *NR3C2* | CM040306.1 | 59433268 | 59433334 |
| *NR3C2* | CM040306.1 | 59434509 | 59434517 |
| *OXTR* | CM040317.1 | 20203310 | 20204344 |
| *POMC* | JADDUC020000174.1 | 259 | 552 |

**S3 Table. Summary metrics for target-enriched enzymatic methyl sequencing (TEEM-Seq).** TEEM-Seq alignments were performed on target region putative promoters and exons to generate mean coverage plots (Fig 2). Since some exon spans were too short for proper alignments, reads were also aligned separately with neighboring intron sequence for proper alignments and then checked for exon depth (see Fig 2d-f). There were 106,178 total bases in the target sequences.

| Individual | Raw reads | Target region mapping efficiency (%) | Target region unique paired-end alignments | Target region sequences after deduplication | Target region mapped reads after deduplication | Mean read depth across putative promoter target regions |
|---|---|---|---|---|---|---|
| BB-17532 | 14,423,547 | 13.7 | 1,965,052 | 124,628 | 249,256 | 318.54x |
| BB-17501 | 10,238,150 | 14.3 | 1,459,928 | 36,825 | 73,650 | 94.44x |
| BB-17455 | 12,105,745 | 16.4 | 1,981,679 | 98,146 | 196,292 | 254.84x |
| BB-17411 | 12,538,149 | 15.5 | 1,934,046 | 68,158 | 136,316 | 178.43x |
| BB-17168 | 5,905,164 | 15.6 | 917,093 | 25,943 | 51,886 | 65.73x |
| BB-14232 | 2,777,787 | 15.9 | 440,740 | 20,203 | 40,406 | 51.6x |
| B-40881 | 37,427,806 | 16.7 | 6,209,452 | 309,228 | 618,456 | 772.53x |

**S4 Table. Summary metrics for whole-genome enzymatic methyl sequencing (WGEM-Seq).** WGEM-Seq alignments were performed on target region putative promoters and exons to generate mean coverage plots (Fig 2). Since some exon spans were too short for proper alignments, reads were also aligned separately with neighboring intron sequence for proper alignments and then checked for exon depth (see Fig 2d-f). There were 106,178 total bases in the target sequences.

| Individual | Raw reads | Full genome mapping efficiency (%) | Full genome unique paired-end alignments | Full genome sequences after deduplication | Mapped reads after deduplication | Mean read depth across genome | Mean read depth across putative promoter target regions |
|---|---|---|---|---|---|---|---|
| BB-17501 | 122,075,233 | 73.8 | 89,559,902 | 75,747,783 | 151,495,566 | 23.07x | 16.04x |
| BB-17168 | 134,336,464 | 73.2 | 97,542,032 | 80,729,566 | 161,459,132 | 26.03x | 17.71x |
| BB-14232 | 92,367,134 | 75.5 | 69,224,635 | 58,745,681 | 117,491,362 | 18.04x | 12.29x |

#### S5 Table. Summary metrics for reduced-representation bisulfite sequencing (RRBS).

| Individual | Raw reads | Full genome mapping efficiency | Full genome unique single-end alignments | Mean CpG coverage across genome* | Mean CpG coverage across putative promoter target regions** |
|---|---|---|---|---|---|
| BB-17532 | 19,253,809 | 52.1 | 9,437,245 | 18.05x | 14.78x |
| BB-17501 | 19,634,960 | 50.1 | 9,504,039 | 18.14x | 14.58x |
| BB-17455 | 14,016,434 | 53.5 | 6,621,698 | 12.80x | 9.51x |
| BB-17411 | 17,805,945 | 55.4 | 7,137,598 | 13.05x | 9.92x |
| BB-17168 | 16,571,855 | 53.8 | 8,649,111 | 16.81x | 13.54x |
| BB-14232 | 19,668,309 | 51.5 | 9,932,118 | 19.40x | 16.03x |
| B-40881 | 29,514,286 | 45.0 | 12,924,898 | 22.87x | 19.18x |

* At 5x coverage or above to avoid low-representation sites in RRBS library.
** Averaging the total count of unmethylated and methylated cytosines for all sites.

**S6 Table. Comparison of target-enriched enzymatic methyl sequencing (TEEM-Seq) and reduced-representation bisulfite sequencing (RRBS) in putative promoter regions.** Libraries for seven of the same individual superb starlings (B-40881, BB-17168, BB-17411, BB-17455, BB-17501, BB-17532, and BB-14232) were sequenced using both methods. The number of shared CpG sites at 5x coverage or above and the mean DNA methylation levels from both methods are shown for each starling sample. Some target regions had very sparse RRBS coverage, resulting in low numbers of shared CpG sites (e.g., *ESRI*, *GNRH1*, *GNRH2*, *MC4R*, and *SERPINA1*). See S2 Table for the length of the putative promoter region targeted for each gene.

| Target region | B-40881 | | | BB-17168 | | | BB-17411 | | | BB-17455 | | | BB-17501 | | | BB-17532 | | | BB-14232 | | |
|---|---|---|---|---|---|---|---|---|---|---|---|---|---|---|---|---|---|---|---|---|---|
| | Number shared CpGs | TEEM-Seq mean methyl | RRBS mean methyl | Number shared CpGs | TEEM-Seq mean methyl | RRBS mean methyl | Number shared CpGs | TEEM-Seq mean methyl | RRBS mean methyl | Number shared CpGs | TEEM-Seq mean methyl | RRBS mean methyl | Number shared CpGs | TEEM-Seq mean methyl | RRBS mean methyl | Number shared CpGs | TEEM-Seq mean methyl | RRBS mean methyl | Number shared CpGs | TEEM-Seq mean methyl | RRBS mean methyl |
| *AR* | 310 | 1.91 | 2.57 | 207 | 3.76 | 2.87 | 253 | 2.48 | 2.43 | 253 | 2.28 | 3.15 | 221 | 3.15 | 3.17 | 270 | 2.14 | 2.37 | 173 | 4.85 | 3.40 |
| *AVPR1A* | 76 | 1.37 | 1.66 | 76 | 2.19 | 1.39 | 76 | 1.16 | 1.27 | 75 | 1.49 | 1.21 | 76 | 1.60 | 2.28 | 76 | 0.96 | 1.49 | 76 | 3.52 | 1.42 |
| *AVPR1B* | 40 | 49.76 | 50.02 | 43 | 51.27 | 57.37 | 43 | 50.05 | 52.32 | 39 | 50.04 | 50.83 | 28 | 48.29 | 53.71 | 36 | 45.86 | 50.25 | 42 | 47.86 | 45.95 |
| *CRH* | 239 | 1.68 | 1.98 | 190 | 3.44 | 2.28 | 224 | 1.47 | 1.58 | 213 | 2.04 | 1.50 | 234 | 2.14 | 1.95 | 213 | 1.14 | 1.25 | 205 | 3.93 | 2.01 |
| *DNMT1* | 154 | 30.32 | 29.62 | 86 | 13.42 | 13.71 | 115 | 17.51 | 16.04 | 132 | 28.58 | 29.53 | 123 | 25.39 | 24.40 | 120 | 27.82 | 28.16 | 72 | 17.35 | 16.41 |
| *DNMT3B* | 76 | 17.53 | 14.97 | 58 | 10.75 | 6.32 | 58 | 10.18 | 13.40 | 62 | 13.81 | 12.50 | 58 | 14.42 | 13.85 | 70 | 12.73 | 13.47 | 49 | 12.60 | 11.39 |
| *EGR1* | 312 | 0.11 | 0.73 | 304 | 1.55 | 0.68 | 290 | 0.60 | 0.91 | 251 | 0.45 | 0.78 | 283 | 0.33 | 0.75 | 293 | 0.12 | 0.73 | 272 | 1.74 | 0.42 |
| *ESR1* | 0 | - | - | 0 | - | - | 0 | - | - | 0 | - | - | 0 | - | - | 0 | - | - | 0 | - | - |
| *FKBP5* | 6 | 70.61 | 61.41 | 6 | 66.19 | 63.89 | 6 | 66.74 | 66.67 | 6 | 70.27 | 67.05 | 6 | 72.46 | 81.40 | 6 | 71.81 | 67.81 | 6 | 66.18 | 62.88 |
| *GNIH* | 54 | 16.58 | 17.89 | 53 | 15.31 | 10.82 | 51 | 17.19 | 16.88 | 53 | 22.10 | 18.58 | 54 | 17.06 | 14.95 | 50 | 10.37 | 12.60 | 50 | 12.02 | 11.74 |
| *GNRH1* | 0 | - | - | 0 | - | - | 1 | 34.62 | - | 1 | 31.18 | - | 0 | - | - | 0 | - | - | 1 | 25.00 | - |
| *GNRHR2 r1*[#] | 8 | 51.54 | 57.44 | 0 | - | - | 6 | 50.37 | 46.53 | 7 | 54.57 | 44.29 | 8 | 53.76 | 43.45 | 8 | 47.92 | 47.80 | 7 | 48.68 | 59.05 |
| *GNRHR2 r2*[#] | 214 | 41.21 | 41.03 | 185 | 54.39 | 52.21 | 176 | 48.86 | 48.33 | 182 | 48.38 | 47.17 | 200 | 50.17 | 48.59 | 209 | 46.42 | 47.16 | 159 | 52.02 | 50.54 |
| *MC2R* | 161 | 0.67 | 1.25 | 151 | 1.83 | 0.45 | 158 | 0.53 | 0.75 | 161 | 0.79 | 0.95 | 150 | 0.53 | 0.69 | 173 | 0.61 | 0.56 | 146 | 3.78 | 0.39 |
| *MC4R* | 0 | - | - | 0 | - | - | 0 | - | - | 0 | - | - | 0 | - | - | 0 | - | - | 0 | - | - |
| *NR3C1* | 219 | 0.26 | 0.39 | 152 | 1.35 | 0.30 | 195 | 0.24 | 0.29 | 179 | 0.16 | 0.47 | 195 | 0.49 | 0.39 | 220 | 0.17 | 0.36 | 172 | 4.13 | 0.24 |
| *NR3C2* | 486 | 0.32 | 0.33 | 355 | 2.17 | 0.70 | 401 | 0.55 | 0.69 | 412 | 0.58 | 0.70 | 412 | 0.85 | 0.75 | 479 | 0.29 | 0.42 | 363 | 4.44 | 0.47 |
| *OXTR* | 145 | 4.32 | 4.50 | 95 | 9.54 | 3.85 | 120 | 4.38 | 3.82 | 146 | 4.36 | 4.17 | 122 | 6.57 | 5.25 | 125 | 3.55 | 3.19 | 101 | 11.48 | 4.39 |
| *POMC* | 23 | 95.04 | 93.73 | 32 | 76.85 | 82.15 | 39 | 91.95 | 93.59 | 25 | 94.13 | 93.04 | 23 | 96.12 | 93.62 | 33 | 95.08 | 94.75 | 18 | 97.13 | 96.35 |
| *SERPINA1* | 0 | - | - | 0 | - | - | 0 | - | - | 0 | - | - | 0 | - | - | 0 | - | - | 0 | - | - |
| *VT* | 10 | 46.16 | 41.38 | 9 | 55.73 | 46.01 | 10 | 45.33 | 45.74 | 10 | 51.09 | 36.63 | 10 | 48.30 | 48.12 | 9 | 46.66 | 50.84 | 9 | 54.80 | 45.34 |
| *VTG1* | 26 | 8.51 | 6.48 | 21 | 22.95 | 18.80 | 26 | 7.78 | 6.50 | 26 | 7.02 | 2.16 | 22 | 15.48 | 18.03 | 26 | 8.46 | 6.35 | 21 | 8.95 | 11.81 |

[#] Two separate gene regions (indicated as r1 and r2) on chromosome 10 with similarity to *GNRHR2*.

**S7 Table. Comparison of whole-genome enzymatic methyl sequencing (WGEM-Seq) and target-enriched enzymatic methyl sequencing (TEEM-Seq) in putative promoter regions.** Libraries for three of the same individual superb starlings (BB-17168, BB-17501, and BB-14232) were sequenced using both methods. The number of shared CpG sites at 5x coverage or above and the mean DNA methylation levels from both methods are shown for each starling sample. Asterisks indicate genes for which more or less than 4000 bp of putative promoter regions were targeted.

| Target region | Putative promoter region length (bp) | BB-17168 | | | BB-17501 | | | BB-14232 | | |
|---|---|---|---|---|---|---|---|---|---|---|
| | | Number shared CpGs | WGEM-Seq mean methyl | TEEM-Seq mean methyl | Number shared CpGs | WGEM-Seq mean methyl | TEEM-Seq mean methyl | Number shared CpGs | WGEM-Seq mean methyl | TEEM-Seq mean methyl |
| *AR* | 4000 | 212 | 9.45 | 8.93 | 226 | 10.27 | 9.22 | 117 | 13.87 | 12.08 |
| *AVPR1A* | 4000 | 60 | 31.17 | 32.61 | 104 | 23.29 | 22.19 | 48 | 18.49 | 17.91 |
| *AVPR1B* | 4000 | 83 | 48.45 | 47.55 | 67 | 40.56 | 41.99 | 53 | 41.89 | 41.02 |
| *CRH* | 4000 | 251 | 7.08 | 5.98 | 239 | 4.58 | 4.55 | 187 | 10.21 | 7.78 |
| *DNMT1* | 4000 | 105 | 40.79 | 41.23 | 131 | 51.64 | 51.07 | 35 | 81.18 | 84.53 |
| *DNMT3A* | 4000 | 288 | 80.64 | 80.67 | 329 | 81.09 | 80.24 | 177 | 83.52 | 81.95 |
| *DNMT3B* | 4000 | 149 | 26.39 | 25.44 | 125 | 33.51 | 34.59 | 98 | 27.97 | 25.48 |
| *EGR1* | 4000 | 328 | 5.47 | 4.53 | 290 | 3.93 | 3.91 | 155 | 7.11 | 5.90 |
| *ESR1* | 4000 | 48 | 55.68 | 58.37 | 44 | 63.91 | 63.63 | 23 | 59.84 | 55.07 |
| *FKBP5* | 4000 | 77 | 40.30 | 39.90 | 78 | 47.61 | 45.69 | 46 | 41.99 | 40.16 |
| *GNIH* | 4000 | 108 | 29.63 | 28.92 | 109 | 28.57 | 29.14 | 71 | 27.31 | 30.54 |
| *GNRH1* | 5726* | 16 | 40.23 | 42.53 | 18 | 45.00 | 47.01 | 4 | 39.40 | 41.60 |
| *GNRHR2 r1*[#] | 4000 | 70 | 46.24 | 48.28 | 66 | 53.62 | 49.83 | 50 | 46.46 | 43.96 |
| *GNRHR2 r2*[#] | 4000 | 189 | 62.80 | 63.77 | 196 | 60.59 | 60.48 | 112 | 58.82 | 58.07 |
| *MC2R* | 3856 | 194 | 10.18 | 7.91 | 143 | 8.57 | 9.21 | 39 | 17.58 | 18.53 |
| *MC4R* | 3948* | 26 | 38.59 | 42.11 | 28 | 39.68 | 40.77 | 21 | 36.13 | 44.64 |
| *NR3C1* | 4000 | 108 | 0.55 | 0.52 | 172 | 0.56 | 0.43 | 141 | 2.11 | 1.56 |
| *NR3C2* | 4000 | 194 | 8.18 | 8.38 | 176 | 6.79 | 8.43 | 33 | 8.24 | 5.76 |
| *OXTR* | 4000 | 103 | 21.72 | 23.73 | 116 | 22.02 | 22.64 | 69 | 29.88 | 28.45 |
| *POMC* | 258** | 20 | 81.25 | 81.14 | 9 | 94.71 | 94.36 | 4 | 100.00 | 99.17 |
| *SERPINA1* | 4000 | 13 | 55.74 | 55.60 | 12 | 44.55 | 46.99 | 12 | 46.83 | 46.46 |
| *VT* | 4000 | 54 | 48.51 | 45.14 | 79 | 44.55 | 41.10 | 78 | 29.51 | 34.43 |
| *VTG1* | 697** | 37 | 25.20 | 23.54 | 13 | 20.00 | 24.22 | 15 | 24.25 | 12.65 |

[*] Since there is no existing annotation in the superb starling reference genome for this gene, we used the zebra finch sequence alignment to target 2 kb upstream in the putative promoter region and 2 kb in the gene body.
[**] Limited sequence upstream of gene.
[#] Two separate gene regions (indicated as r1 and r2) on chromosome 10 with similarity to *GNRHR2*.

**S8 Table. Comparison of samples run as duplicate or triplicate libraries using target-enriched enzymatic methyl sequencing (TEEM-Seq).** Samples for four of the same individual superb starlings (BB-17411, BB-17455, B-40881, and BB-17532) were made into duplicate (or triplicate for BB-17532) libraries and sequenced. The number of shared CpG sites at 5x coverage or above and the mean DNA methylation levels from each library are shown for each starling sample. Asterisks indicate genes for which more or less than 4000 bp of putative promoter regions were targeted.

| Target region | Putative promoter region length (bp) | BB-17411 | | | BB-17455 | | | B-40881 | | | BB-17532 | | | | | | | | |
|---|---|---|---|---|---|---|---|---|---|---|---|---|---|---|---|---|---|---|---|
| | | Number shared CpGs | Library 1 mean methyl | Library 2 mean methyl | Number shared CpGs | Library 1 mean methyl | Library 2 mean methyl | Number shared CpGs | Library 1 mean methyl | Library 2 mean methyl | Number shared CpGs | Library 1 mean methyl | Library 2 mean methyl | Number shared CpGs | Library 1 mean methyl | Library 3 mean methyl | Number shared CpGs | Library 2 mean methyl | Library 3 mean methyl |
| *AR* | 4000 | 430 | 5.10 | 5.18 | 431 | 4.72 | 5.09 | 444 | 4.63 | 4.65 | 377 | 5.29 | 5.28 | 325 | 5.95 | 5.94 | 312 | 6.09 | 6.19 |
| *AVPR1A* | 4000 | 152 | 16.39 | 16.26 | 151 | 16.06 | 15.84 | 154 | 16.58 | 15.93 | 152 | 15.18 | 14.27 | 150 | 15.16 | 15.21 | 150 | 14.09 | 15.21 |
| *AVPR1B* | 4000 | 125 | 45.47 | 44.64 | 123 | 45.04 | 46.47 | 124 | 43.77 | 44.55 | 123 | 42.03 | 41.49 | 117 | 42.80 | 40.31 | 117 | 41.93 | 40.31 |
| *CRH* | 4000 | 364 | 4.37 | 4.27 | 370 | 5.18 | 5.71 | 372 | 4.57 | 4.86 | 357 | 4.24 | 4.18 | 334 | 4.35 | 5.05 | 333 | 4.23 | 5.07 |
| *DNMT1* | 4000 | 286 | 50.07 | 48.21 | 288 | 47.54 | 47.37 | 301 | 46.90 | 46.72 | 247 | 55.90 | 54.81 | 211 | 61.23 | 60.66 | 211 | 60.18 | 60.66 |
| *DNMT3A* | 4000 | 439 | 78.10 | 77.33 | 473 | 77.95 | 76.62 | 513 | 79.37 | 79.64 | 418 | 79.40 | 79.90 | 366 | 79.26 | 78.25 | 353 | 79.68 | 78.06 |
| *DNMT3B* | 4000 | 183 | 25.19 | 25.21 | 183 | 25.60 | 25.24 | 183 | 26.60 | 26.26 | 183 | 23.03 | 23.25 | 183 | 23.03 | 23.61 | 183 | 23.25 | 23.61 |
| *EGR1* | 4000 | 456 | 2.63 | 2.52 | 458 | 2.84 | 2.55 | 472 | 2.41 | 2.62 | 459 | 2.31 | 2.59 | 415 | 2.53 | 3.72 | 415 | 2.83 | 3.72 |
| *ESR1* | 4000 | 48 | 58.81 | 55.23 | 47 | 59.74 | 62.25 | 55 | 57.44 | 56.77 | 50 | 59.39 | 61.09 | 44 | 59.18 | 61.74 | 44 | 61.23 | 61.74 |
| *FKBP5* | 4000 | 103 | 40.30 | 38.22 | 103 | 43.50 | 43.20 | 104 | 41.62 | 41.68 | 103 | 39.88 | 37.65 | 102 | 40.27 | 40.59 | 102 | 38.02 | 40.59 |
| *GNIH* | 4000 | 129 | 28.68 | 26.61 | 128 | 31.30 | 34.05 | 131 | 28.15 | 28.34 | 129 | 23.85 | 24.07 | 127 | 23.91 | 23.14 | 127 | 24.01 | 23.14 |
| *GNRH1* | 5726* | 22 | 45.52 | 48.92 | 22 | 41.15 | 41.98 | 24 | 42.94 | 42.50 | 23 | 40.31 | 38.39 | 21 | 41.18 | 43.82 | 21 | 39.23 | 43.82 |
| *GNRHR2 (r1)*# | 4000 | 85 | 47.92 | 48.84 | 83 | 47.68 | 48.99 | 89 | 44.75 | 44.73 | 84 | 45.35 | 44.93 | 82 | 46.45 | 45.93 | 82 | 46.02 | 45.93 |
| *GNRHR2 (r2)*# | 4000 | 388 | 51.86 | 49.94 | 371 | 50.45 | 50.86 | 389 | 48.30 | 48.91 | 370 | 49.85 | 51.30 | 327 | 54.42 | 55.53 | 321 | 55.60 | 55.63 |
| *MC2R* | 3856 | 279 | 4.62 | 4.97 | 279 | 4.81 | 5.01 | 293 | 5.09 | 5.04 | 279 | 4.59 | 5.71 | 279 | 4.59 | 4.54 | 272 | 5.86 | 4.65 |
| *MC4R* | 3948* | 32 | 42.24 | 38.52 | 31 | 41.43 | 40.61 | 32 | 43.63 | 41.79 | 32 | 41.10 | 39.70 | 32 | 41.10 | 37.45 | 32 | 39.70 | 37.45 |
| *NR3C1* | 4000 | 333 | 0.43 | 0.30 | 314 | 0.31 | 0.22 | 392 | 0.22 | 0.25 | 316 | 0.20 | 0.25 | 210 | 0.27 | 0.16 | 206 | 0.17 | 0.06 |
| *NR3C2* | 4000 | 756 | 2.21 | 2.21 | 752 | 2.35 | 2.05 | 824 | 1.96 | 1.91 | 719 | 1.90 | 2.45 | 589 | 2.27 | 2.77 | 560 | 2.87 | 2.70 |
| *OXTR* | 4000 | 228 | 15.91 | 17.68 | 204 | 18.45 | 19.51 | 244 | 15.80 | 16.57 | 205 | 16.34 | 17.21 | 180 | 17.96 | 20.64 | 197 | 17.20 | 18.96 |
| *POMC* | 258** | 32 | 92.09 | 92.61 | 23 | 93.37 | 94.52 | 48 | 93.50 | 93.16 | 42 | 94.47 | 91.35 | 34 | 95.19 | 93.79 | 31 | 90.65 | 93.19 |
| *SERPINA1* | 4000 | 15 | 44.88 | 45.92 | 13 | 53.40 | 47.63 | 16 | 47.16 | 46.39 | 15 | 46.63 | 42.52 | 14 | 46.69 | 41.69 | 14 | 42.70 | 41.69 |
| *VT* | 4000 | 133 | 37.30 | 35.46 | 133 | 38.46 | 40.57 | 137 | 37.73 | 37.80 | 130 | 34.25 | 34.27 | 128 | 34.43 | 33.80 | 126 | 34.40 | 33.92 |
| *VTG1* | 697** | 56 | 9.77 | 7.91 | 55 | 8.11 | 8.46 | 70 | 7.23 | 7.64 | 55 | 8.66 | 10.08 | 40 | 10.24 | 9.74 | 40 | 13.51 | 9.74 |

* Since there is no existing annotation in the superb starling reference genome for this gene, we used the zebra finch sequence alignment to target 2 kb upstream in the putative promoter region and 2 kb in the gene body.
** Limited sequence upstream of gene.
### Two separate gene regions (indicated as r1 and r2) on chromosome 10 with similarity to *GNRHR2*.

**S9 Table. Comparison of whole-genome enzymatic methyl sequencing (WGEM-Seq) and target-enriched enzymatic methyl sequencing (TEEM-Seq) in exonic regions.** Libraries for three of the same individual superb starlings (BB-17168, BB-17501, and BB-14232) were sequenced using both methods. The number of shared CpG sites at 5x coverage or above and the mean DNA methylation levels from both methods are shown for each starling sample.

| Target region | BB-17168 | | | BB-17501 | | | BB-14232 | | |
|---|---|---|---|---|---|---|---|---|---|
| | **Number shared CpGs** | **WGEM-Seq mean methyl** | **TEEM-Seq mean methyl** | **Number shared CpGs** | **WGEM-Seq mean methyl** | **TEEM-Seq mean methyl** | **Number shared CpGs** | **WGEM-Seq mean methyl** | **TEEM-Seq mean methyl** |
| *AR* | 151 | 32.45 | 31.65 | 163 | 36.96 | 37.46 | 103 | 40.21 | 40.34 |
| *AVPR1A* | 229 | 8.42 | 7.06 | 172 | 9.03 | 8.21 | 115 | 9.19 | 10.20 |
| *AVPR1B* | 111 | 66.61 | 68.59 | 116 | 64.81 | 64.81 | 72 | 80.31 | 76.70 |
| *CRH* | 41 | 2.32 | 0.69 | 40 | 0.00 | 0.17 | 15 | 6.41 | 3.31 |
| *EGR1* | 155 | 21.54 | 20.53 | 158 | 25.03 | 25.06 | 120 | 25.57 | 25.76 |
| *FKBP5* | 66 | 64.57 | 63.02 | 60 | 67.84 | 64.85 | 31 | 68.63 | 68.82 |
| *GNRHR2 r1*# | 53 | 70.84 | 70.77 | 62 | 72.68 | 73.50 | 39 | 68.13 | 68.07 |
| *GNRHR2 r2*# | 120 | 76.15 | 76.49 | 130 | 78.31 | 78.39 | 89 | 78.93 | 82.03 |
| *NR3C1* | 260 | 57.74 | 58.36 | 241 | 57.52 | 60.04 | 119 | 47.73 | 51.44 |
| *NR3C2* | 145 | 85.30 | 82.91 | 141 | 85.63 | 85.53 | 84 | 83.30 | 82.98 |
| *OXTR* | 65 | 55.26 | 56.32 | 58 | 69.73 | 67.39 | 42 | 55.55 | 60.35 |
| *POMC* | 22 | 88.21 | 88.18 | 26 | 86.28 | 87.31 | 21 | 92.18 | 90.93 |

### Two separate gene regions (indicated as r1 and r2) on chromosome 10 with similarity to *GNRHR2*.

**S10 Table. Comparison of whole-genome enzymatic methyl sequencing (WGEM-Seq) and reduced-representation bisulfite sequencing (RRBS) in random 2.5 kb "promoter" regions.** WGEM-Seq and RRBS data were compared in 20 random 2.5 kb "promoter" regions that were not included in the bait set because we expected that some RRBS regions corresponding to target ranges might have sparse coverage in comparison to the high coverage bait alignments. The position of each random 2.5 kb "promoter" region in the superb starling genome (Lasu_v2), the chromosome number, and "promoter" orientation ('rev' indicates 'reversed'). Libraries for three of the same individual superb starlings (BB-17168, BB-17501, and BB-14232) were sequenced using both methods. The sequence start and ending locations, the number of shared CpG sites at 5x coverage or above, and the mean DNA methylation levels from both methods are shown for each starling sample.

| Genome position | BB-17168 | | | | | | BB-17501 | | | | | | BB-14232 | | | | | |
|---|---|---|---|---|---|---|---|---|---|---|---|---|---|---|---|---|---|---|
| *Chromosome* | **Start sequence location (1-based)** | **End sequence location** | **Length (bp)** | **Number shared CpGs** | **WGEM-Seq mean methyl** | **RRBS mean methyl** | **Start sequence location (1-based)** | **End sequence location** | **Length (bp)** | **Number shared CpGs** | **WGEM-Seq mean methyl** | **RRBS mean methyl** | **Start sequence location (1-based)** | **End sequence location** | **Length (bp)** | **Number shared CpGs** | **WGEM-Seq mean methyl** | **RRBS mean methyl** |
| CM040302.1 *chr 1* | 33752942 | 33753769 | 828 | 62 | 0.55 | 0.42 | 33752942 | 33753751 | 810 | 70 | 0.00 | 0.27 | 33753270 | 33753768 | 499 | 59 | 2.67 | 0.53 |
| CM040302.1 *chr 1 (rev)* | 62786630 | 62787143 | 514 | 28 | 1.00 | 0.25 | 62786427 | 62787143 | 717 | 55 | 0.61 | 0.16 | 62786638 | 62787130 | 493 | 29 | 4.52 | 0.13 |
| CM040303.1 *chr 1A* | 27626923 | 27627048 | 126 | 14 | 0.00 | 0.79 | 27626860 | 27627068 | 209 | 21 | 0.00 | 0.40 | 27626922 | 27627048 | 127 | 0 | N/A | N/A |
| CM040303.1 *chr 1A (rev) r1*[#] | 10576313 | 10577590 | 1278 | 59 | 2.22 | 3.19 | 10576294 | 10577591 | 1298 | 52 | 0.00 | 1.84 | 10576289 | 10577633 | 1345 | 88 | 1.84 | 0.69 |
| CM040303.1 *chr 1A (rev) r2*[#] | 39265876 | 39267417 | 1542 | 134 | 0.25 | 0.28 | 39265876 | 39267417 | 1542 | 219 | 0.00 | 0.40 | 39265911 | 39267426 | 1516 | 177 | 4.38 | 0.24 |
| CM040305.1 *chr 3 r1*[#] | 12188281 | 12189069 | 789 | 37 | 2.05 | 1.19 | 12188282 | 12189069 | 788 | 26 | 1.28 | 2.21 | 12188282 | 12189082 | 801 | 20 | 10.33 | 3.36 |
| CM040305.1 *chr 3 r2*[#] | 43713632 | 43713963 | 332 | 37 | 4.11 | 0.12 | 43713632 | 43713963 | 332 | 21 | 0.00 | 0.25 | 43713663 | 43713963 | 301 | 7 | 0.00 | 1.02 |
| CM040306.1 *chr 4 (rev)* | 18836665 | 18837815 | 1151 | 71 | 5.04 | 0.83 | 18836672 | 18837050 | 379 | 13 | 0.00 | 0.00 | 18836881 | 18837814 | 934 | 62 | 7.37 | 1.44 |
| CM040307.1 *chr 4A (rev)* | 9285901 | 9287354 | 1454 | 60 | 3.89 | 4.33 | 9285901 | 9287355 | 1455 | 34 | 8.50 | 6.42 | 9287200 | 9287368 | 169 | 29 | 2.53 | 0.39 |
| CM040308.1 *chr 5A* | 30585721 | 30585808 | 88 | 10 | 2.43 | 13.00 | 30585691 | 30585808 | 118 | 19 | 24.78 | 23.82 | 30585707 | 30585809 | 103 | 21 | 10.98 | 2.73 |
| CM040310.1 *chr 6A* | 12600601 | 12601812 | 1212 | 10 | 28.44 | 25.18 | 12600601 | 12601812 | 1212 | 9 | 35.41 | 31.13 | 12601734 | 12601811 | 78 | 4 | 2.08 | 1.92 |
| CM040312.1 *chr 7 (rev)* | 26600657 | 26601613 | 957 | 54 | 2.31 | 1.71 | 26600729 | 26601612 | 884 | 79 | 0.75 | 0.86 | 26600998 | 26601403 | 406 | 60 | 6.17 | 0.14 |
| CM040317.1 *chr 12* | 518243 | 518850 | 608 | 32 | 2.41 | 1.42 | 518244 | 518926 | 683 | 13 | 0.00 | 0.96 | 518251 | 518809 | 559 | 20 | 11.24 | 0.66 |
| CM040322.1 *chr 18 (rev)* | 11611593 | 11612416 | 824 | 65 | 11.80 | 11.72 | 11611593 | 11612417 | 825 | 47 | 12.75 | 10.72 | 11611593 | 11612429 | 837 | 69 | 6.34 | 4.75 |
| CM040325.1 *chr 21 (rev)* | 641318 | 641744 | 427 | 13 | 4.18 | 0.59 | 641317 | 641505 | 189 | 24 | 0.83 | 0.21 | 641314 | 641506 | 193 | 32 | 7.50 | 0.63 |
| CM040327.1 *chr 23* | 7414348 | 7415383 | 1036 | 24 | 0.83 | 0.00 | 7414415 | 7415713 | 1299 | 14 | 1.43 | 1.43 | 7414918 | 7415028 | 111 | 25 | 1.60 | 0.30 |
| CM040329.1 *chr 25* | 3307103 | 3308528 | 1426 | 52 | 9.76 | 7.90 | 3307138 | 3308528 | 1391 | 57 | 14.58 | 13.62 | 3307852 | 3308095 | 244 | 28 | 3.37 | 0.11 |
| CM040330.1 *chr 26 (rev)* | 1656448 | 1657477 | 1030 | 41 | 13.25 | 11.71 | 1656887 | 1657532 | 646 | 19 | 39.37 | 30.22 | 1656256 | 1657478 | 1223 | 49 | 11.89 | 5.13 |
| CM040333.1 *chr 29* | 2390588 | 2392092 | 1505 | 35 | 0.00 | 0.00 | 2390589 | 2392092 | 1504 | 39 | 0.51 | 0.52 | 2390750 | 2390868 | 119 | 5 | 0.00 | 0.00 |
| CM040335.1 *chr Z* | 20569346 | 20569391 | 46 | 9 | 0.00 | 1.85 | 20568984 | 20569391 | 408 | 34 | 0.42 | 0.35 | 20569345 | 20569370 | 26 | 7 | 4.76 | 0.00 |

[#] Two separate gene regions (indicated as r1 and r2) on chromosome 1A (rev) and chromosome 3.

**S11 Table. Comparison of whole-genome enzymatic methyl sequencing (WGEM-Seq) and target-enriched enzymatic methyl sequencing (TEEM-Seq) in putative promoter regions for TEEM-Seq data without deduplication.** Libraries for three of the same individual superb starlings (BB-17168, BB-17501, and BB-14232) were sequenced using both methods. The number of shared CpG sites at 5x coverage or above and the mean DNA methylation levels from both methods are shown for each starling sample. WGEM-Seq data is deduplicated by default. S7 Table shows the same comparison for TEEM-Seq data with deduplication. Given the qualitatively similar results using deduplication and non-deduplication of TEEM-Seq data, similar to RRBS analysis, deduplication may not be necessary for TEEM-Seq analysis. Asterisks indicate genes for which more or less than 4000 bp of putative promoter regions were targeted.

| Target region | Putative promoter region length (bp) | BB-17168 | | | BB-17501 | | | BB-14232 | | |
|---|---|---|---|---|---|---|---|---|---|---|
| | | Number shared CpGs | WGEM-Seq mean methyl | TEEM-Seq mean methyl | Number shared CpGs | WGEM-Seq mean methyl | TEEM-Seq mean methyl | Number shared CpGs | WGEM-Seq mean methyl | TEEM-Seq mean methyl |
| *AR* | 4000 | 212 | 9.45 | 8.76 | 226 | 10.27 | 9.30 | 117 | 13.87 | 11.75 |
| *AVPR1A* | 4000 | 60 | 31.17 | 32.96 | 104 | 23.29 | 22.01 | 48 | 18.49 | 18.24 |
| *AVPR1B* | 4000 | 83 | 48.45 | 48.08 | 67 | 40.56 | 41.63 | 53 | 41.89 | 41.99 |
| *CRH* | 4000 | 251 | 7.08 | 6.10 | 239 | 4.58 | 4.68 | 187 | 10.21 | 8.56 |
| *DNMT1* | 4000 | 105 | 40.79 | 42.10 | 131 | 51.64 | 50.57 | 35 | 81.18 | 83.86 |
| *DNMT3A* | 4000 | 288 | 80.64 | 80.78 | 329 | 81.09 | 80.08 | 177 | 83.52 | 82.58 |
| *DNMT3B* | 4000 | 149 | 26.39 | 25.96 | 125 | 33.51 | 34.84 | 98 | 27.97 | 24.37 |
| *EGR1* | 4000 | 328 | 5.47 | 4.78 | 290 | 3.93 | 3.84 | 155 | 7.11 | 6.14 |
| *ESR1* | 4000 | 48 | 55.68 | 58.61 | 44 | 63.91 | 62.57 | 23 | 59.84 | 55.63 |
| *FKBP5* | 4000 | 77 | 40.30 | 39.85 | 78 | 47.61 | 45.59 | 46 | 41.99 | 39.25 |
| *GNIH* | 4000 | 108 | 29.63 | 27.98 | 109 | 28.57 | 28.95 | 71 | 27.31 | 31.62 |
| *GNRH1* | 5726* | 16 | 40.23 | 42.88 | 18 | 45.00 | 50.10 | 4 | 39.40 | 37.41 |
| *GNRHR2 r1*# | 4000 | 70 | 46.24 | 47.89 | 66 | 53.62 | 49.85 | 50 | 46.46 | 43.08 |
| *GNRHR2 r2*# | 4000 | 189 | 62.80 | 64.01 | 196 | 60.59 | 60.38 | 112 | 58.82 | 57.18 |
| *MC2R* | 3856 | 194 | 10.18 | 7.94 | 143 | 8.57 | 9.37 | 39 | 17.58 | 19.16 |
| *MC4R* | 3948* | 26 | 38.59 | 43.01 | 28 | 39.68 | 39.57 | 21 | 36.13 | 43.00 |
| *NR3C1* | 4000 | 108 | 0.55 | 0.42 | 172 | 0.56 | 0.34 | 141 | 2.11 | 1.39 |
| *NR3C2* | 4000 | 194 | 8.18 | 8.22 | 176 | 6.79 | 8.79 | 33 | 8.24 | 6.44 |
| *OXTR* | 4000 | 103 | 21.72 | 22.96 | 116 | 22.02 | 22.41 | 69 | 29.88 | 28.38 |
| *POMC* | 258** | 20 | 81.25 | 76.16 | 9 | 94.71 | 96.31 | 4 | 100.00 | 98.81 |
| *SERPINA1* | 4000 | 13 | 55.74 | 56.68 | 12 | 44.55 | 43.07 | 12 | 46.83 | 45.02 |
| *VT* | 4000 | 54 | 48.51 | 46.11 | 79 | 44.55 | 40.16 | 78 | 29.51 | 33.89 |
| *VTG1* | 697** | 37 | 25.20 | 19.93 | 13 | 20.00 | 21.50 | 15 | 24.25 | 13.98 |

* Since there is no existing annotation in the superb starling reference genome for this gene, we used the zebra finch sequence alignment to target 2 kb upstream in the putative promoter region and 2 kb in the gene body.

** Limited sequence upstream of gene.

### Two separate gene regions (indicated as r1 and r2) on chromosome 10 with similarity to *GNRHR2*.

**S12 Table. Summary of differences in approach and cost, as well as the pros and cons, of whole-genome enzymatic methyl sequencing (WGEM-Seq), reduced-representation bisulfite sequencing (RRBS), and target-enriched enzymatic methyl sequencing (TEEM-Seq).**

| | WGEM-Seq | RRBS | TEEM-Seq |
|---|---|---|---|
| **Input DNA*** | 80-100 ng | 100 ng | 80-100 ng |
| **Percent CpG sites covered** | 90-95% | 10-15% | user defined |
| **Number samples per reaction** | 1 sample | 1 sample | up to 96 samples |
| **Average sequencing depth per sample** | 27.5 Gb | 1.75 Gb | 2.5 Gb |
| **Preparation time (for 96 samples)** | 4 days | 3 days | 5 days |
| **Per sample costs (as of 2022)** | | | |
| **Reagents**** | $40.00 | $135.00 | $10.00 |
| **Sequencing#** | $275.00 | $17.50 | $25.00 |
| **TOTAL** | $315.00 | $152.50 | $35.00 |
| **Pros** | Targets nearly all CpG sites in genome | Targets ~10-15% of CpG sites in genome<br>Good coverage in putative promoter regions | Near-complete coverage for targeted regions<br>No bias toward CpG rich regions |
| **Cons** | Expensive per sample cost | CpG sites can be missed outside of CpG islands<br>Bisulfite treatment fragments DNA and results in GC bias | Longer prep time<br>High read duplicate rate |

* Based on NEB and NuGEN kits. EM-Seq can theoretically be used down to 10 ng.

** Includes kits (for WGEM-Seq, RRBS, and TEEM-Seq), other reagents (extra beads and probes for TEEM-Seq), sample quality control (for WGEM-Seq, RRBS, and TEEM-Seq), and DNA shearing (for WGEM-Seq and TEEM-Seq). RRBS prices could be reduced using other types of commercial kits or protocols.

### Assuming $10.00 per Gb for 150 PE.
